# Innate immune responses to *Plasmodium falciparum* disrupt the blood-brain barrier

**DOI:** 10.1101/2025.10.20.683440

**Authors:** Alina Batzilla, Pia Dernick, Jon Bezney, Andreas R. Gschwind, Fumio Nakaki, Mireia Altadill, Hannah Fleckenstein, Livia Piatti, Silvia Sanz Sender, Manuel Fiegl, Olawunmi Rashidat Oyerinde, Borja Lopez Gutierrez, Miguel Lopez-Botet, James Sharpe, Lars Steinmetz, Gemma Moncunill, Maria Bernabeu

## Abstract

*Plasmodium falciparum* accumulation at the blood-brain barrier (BBB) is a hallmark of cerebral malaria, a life-threatening complication. Conversely, the contribution of the immune response to vascular injury has long been debated. Here, we studied the role of innate immune cells as potential effectors of vascular damage using a human *in vitro* 3D-BBB model. Parasite-stimulated immune cells from malaria-naïve donors increased adhesion to microvessels, at least partly through LFA-1. This caused barrier disruption and inflammatory activation of BBB cells. Secretion of TNF-α, IFN-γ, and granzyme B by monocytes, NK and γδ T cells correlated with vascular injury, and accumulation of immune cells was required for local barrier damage. Our computational analysis disentangled pathogenic mechanisms driven specifically by either *P. falciparum* parasite or immune cells, as well as shared pathways. These findings demonstrate how vascular-immune interactions may contribute to vascular injury in cerebral malaria and point towards the potential of immunomodulatory therapeutics.

## Introduction

Malaria accounts for more than 600,000 deaths each year.^1^ A substantial fraction are due to cerebral malaria (CM), a life-threatening complication of *Plasmodium falciparum* infection with a fatality rate of 15-20%^2^ and long-term disabilities in 50% of survivors,^3^ even after parenteral administration of effective antimalarials. Despite the urgent need for new adjuvant therapies to alleviate the disease burden, progress is hindered by our limited understanding of the underlying pathogenic mechanisms. CM is characterized by severe brain swelling,^4,5^ as a consequence of blood-brain barrier (BBB) disruption. The sequestration of *P. falciparum-*infected red blood cells (iRBC) in the microvasculature has long been recognized as the main pathogenic event leading to disease.^6,7^ However, recent studies have revealed the accumulation of diverse immune cell types within the cerebral microvasculature of CM patients, including monocytes^8^ and CD8+ T cells.^9,10^ The direct role of the immune response in BBB breakdown, independent of iRBC, remains unclear. Specifically, whether immune processes are causal drivers or a secondary response to vascular injury is largely unknown. The role of the immune response in CM has been primarily studied in the murine experimental cerebral malaria (ECM) *P. berghei* model. There, studies have identified CD8+ T-cells as one of the main effector cells during disease complications,^11^ with microvascular recruitment being mostly driven by chemokines and endothelial activation.^12,13^ Given the fundamental differences between *Plasmodium* species and their host-specific immune responses, we currently lack important knowledge on the contribution of immune cells to human CM pathology. Bioengineered 3D microvascular systems with refined microfluidic networks provide an alternative platform to address this long-standing question. These models, ranging from endothelial-only constructs^14–16^ to 3D-BBB systems, incorporating brain pericytes^17^ and astrocytes,^18^ have successfully recapitulated key aspects of *P. falciparum*-induced pathogenesis.

Here we developed an immunocompetent human *in vitro* 3D-BBB model perfused with immune cells, and performed complex multimodal studies, including single-cell RNA sequencing (scRNA-seq) and functional permeability analysis. Our study aims to recapitulate immune responses upon the first encounters with the parasite, which often result in severe disease. We revealed a pronounced role for parasite-activated leukocytes in BBB breakdown in CM, which occurred in the absence of iRBC. Stimulation of peripheral blood mononuclear cells with *P. falciparum* (*Pf-*PBMC) increased their binding to the 3D-BBB model. This enhancement was particularly pronounced in CD8+ effector and γδ T cells, and occurred in conjunction with switches to lymphocyte function-associated antigen 1 (LFA-1) high-affinity conformation. Adherent leukocytes induced inflammatory pathways in all BBB cell types, and triggered barrier-disruptive endothelial transcriptional programs. These changes were accompanied by an increase in BBB permeability, significantly abrogated by inhibiting leukocyte binding. Integration with previous scRNA-seq data on disruptive mechanisms directly induced by *P. falciparum* revealed shared dysregulated pathways between parasite and innate immune cells, as well as immune-specific pathogenic mechanisms. Our study reveals a prominent, independent contribution of *P. falciparum*-activated leukocytes to BBB breakdown in CM, offering new mechanistic insights and therapeutic opportunities.

## Results

### *P. falciparum-*stimulation induces activation of monocytes, γδ T cells, and NK cells

Blood transcriptome analysis from CM patients has revealed an upregulation of inflammatory, cytotoxic, and general immune cell activation signatures.^19^ First, we decided to characterize the effect of *P. falciparum*-stimulation on PBMC. We stimulated PBMC from malaria-naïve donors overnight with purified, tightly synchronized, late-stage *P. falciparum*-iRBC (parasite line HB3var03) at 37-45 hours post infection, which ensures parasite egress during the stimulation time and the absence of viable parasites in subsequent 3D-BBB perfusion experiments (Figure 1A). Stimulation was done at a ratio of 3:1 iRBC to PBMC, which corresponds to 0.18% parasitemia, a *P. falciparum* concentration relatively low compared to those found in CM patients (0.5 - 5%).^5^ After stimulation of PBMC with *P. falciparum* or with uninfected RBC (referred as control hereafter), we assessed the activation of the different immune cell populations by scRNA-seq and fluorescence-activated cell sorting (FACS). After quality control and integration of datasets from three different PBMC donors, we obtained 26,483 single-cell transcriptomes, which were visualized using uniform manifold approximation and projection (UMAP) (Figures 1B and 1C). When the data were separated by condition, a noticeable transcriptomic shift in immune cell profiles was observed following *P. falciparum*-iRBC stimulation (Figure 1B). We annotated the major PBMC cell types by unsupervised clustering and classical immune cell type markers (Figures 1C and S1A-B). We observed no significant changes in cell type proportions caused by *P. falciparum*-stimulation, indicating no major differences in proliferation or cell death of specific immune cell populations (Figure 1D). Differential gene expression analysis revealed a high number of differentially expressed (DE) genes in monocytes (2358); a moderate number in T cells (1168), NK cells (750), and B cells (584); and a minimal number of DE genes in dendritic cells (29) (Figure 1E).

**Figure 1.**
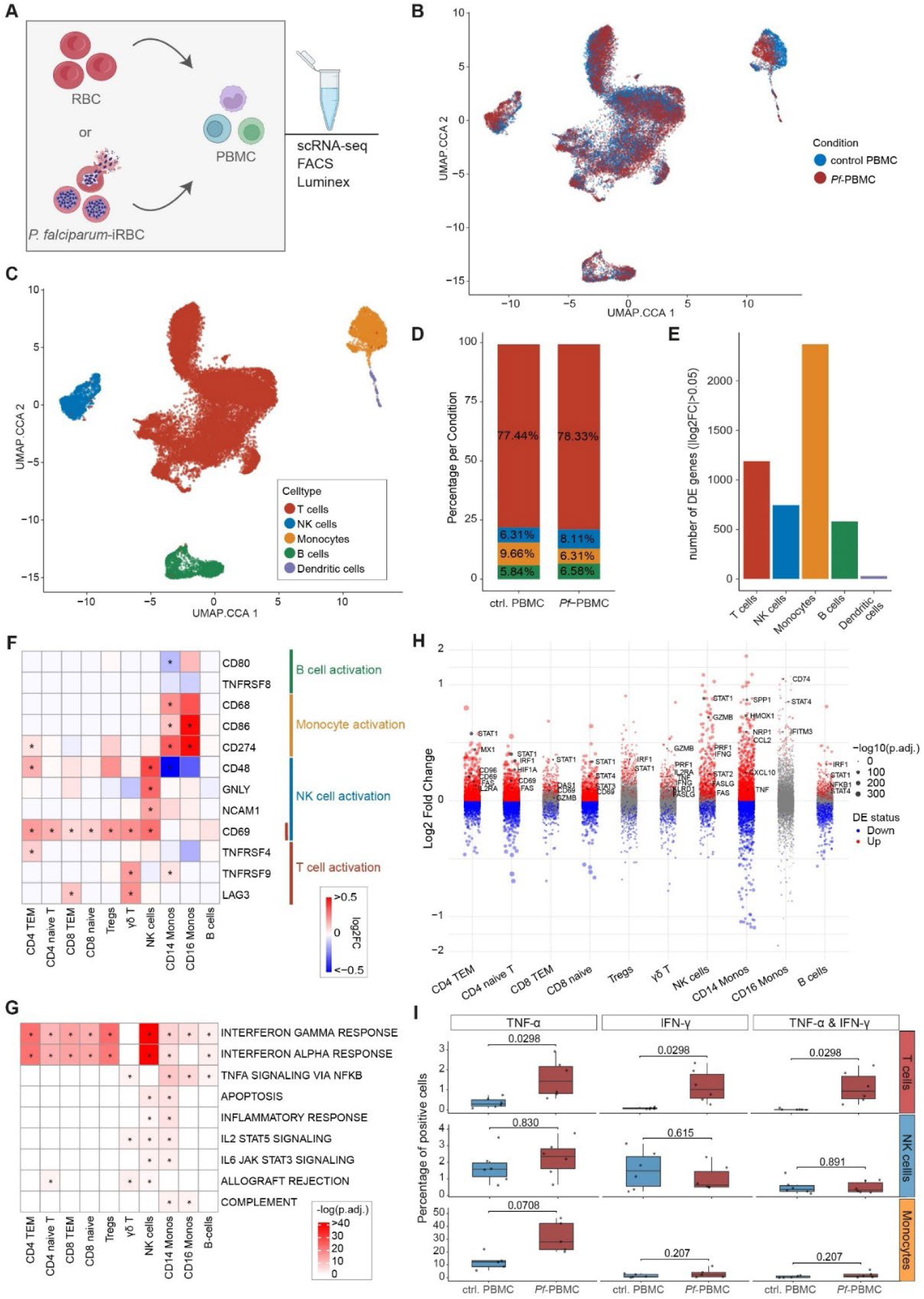
*P. falciparum*-iRBC induce activation of PBMC with increased expression of TNF-α, IFN-γ, and cytotoxic markers. **A.** Schematic experimental overview of PBMC stimulation with *P. falciparum*-iRBC or control RBC and multimodal analysis. **B-C.** UMAP visualization of sequenced PBMC, colored by experimental condition (**B**) and annotated cell type identity (**C**). **D.** Bar plot showing the cell type proportions in control or *Pf*-PBMC. **E.** Bar plot showing the number of significantly differentially expressed (DE) genes (false discovery rate (FDR) < 0.05, |log2FC| > 0.05) in each immune cell type in control or *Pf-*PBMC. **F.** Heatmap of log2-transformed fold change (log2FC) values for selected cell type–specific activation markers, highlighting DE genes (FDR < 0.05, |log2FC| > 0.05) with asterisks. **G.** Gene set over-representation analysis using MSigDB hallmark gene sets on significantly upregulated genes (FDR < 0.05, log2FC > 0.2) across all immune cell types. Displayed gene sets show significant over-representation in at least two cell types. **H.** Volcano plots of DE genes in all immune cell types upon iRBC-stimulation plotting the log2FC and the statistical significance (-log10 of the FDR). Significantly up- or downregulated genes (FDR < 0.05) are marked in red or blue and selected upregulated genes from MSigDB hallmark gene sets are labeled. **I.** Percentage of T cells, NK cells and monocytes expressing TNF-α, IFN-γ or both, assessed by intracellular cytokine staining and FACS. Each dot represents an independent FACS experiment with a different PBMC donor (n = 5–6). Paired Student’s t-test with Benjamini–Hochberg (BH) correction.

Cell type-specific activation markers appeared to be upregulated, with the strongest effect in CD14+ monocytes, followed by NK cells, CD16+ monocytes and T cells. Among T cells, innate-like / γδ T cells were strongly activated, while CD8+ and CD4+ effector memory T cells (CD8+/ CD4+ TEM) showed a modest upregulation of activation markers (Figure 1F). Therefore, the effect was not only constrained to classical innate immune cells but could also be detected in a subset of T cells. FACS analysis showed that activation within the innate-like T cell compartment was primarily confined to γδ T cells, confirming earlier studies (Figure S1C).^20^ *CellChat* ligand-receptor interaction analysis^21^ detected strong signaling originating from classical innate immune cells, CD14+ and CD16+ monocytes and NK cell, potentially contributing to T cell inflammatory activation (Figure S1D), suggesting bystander activation. No upregulation of B cell activation markers was observed (Figure 1F). Using gene set over-representation analysis, we observed a strong induction of several key innate immune response pathways, including interferon (IFN)-γ and IFN-α, tumor necrosis factor (TNF-α) signaling via NF-κB, inflammatory response, and JAK-STAT pathways (Figures 1G and H). Notably, NK cells, γδ T cells, and CD8+ TEM upregulated cytotoxic proteins and cell death regulators like granzyme B, perforin, and Fas ligand (Figure 1G and H). These proteins have been shown to contribute to vascular damage,^22^ with granzyme B found to be increased in blood^23^ and in brain postmortem samples^9^ of CM patients. Moreover, cytokines elevated in CM patients,^24^ such as TNF-α and IFN-γ, were significantly upregulated in monocytes, γδ T cells, and NK cells (Figures 1H and S1C). Intracellular cytokine staining and FACS analysis confirmed increased expression of TNF-α in monocytes and co-expression of both TNF-α and IFN-γ levels in T cells (Figure 1I). Taken together, *P. falciparum-*iRBC stimulation of monocytes, T cells and NK cells induced an increased expression of activation markers, cytokines, and factors involved in immune cell-induced vascular damage.

### *P. falciparum*-stimulated PBMC show increased binding to endothelial cells in 3D microvascular blood-brain barrier model

Leukocyte interaction with endothelial cells is a key step for vascular damage in multiple diseases.^25^ To better understand the interactions of *P. falciparum-*activated leukocytes with microvessels, we developed an immunocompetent *in vitro* 3D-BBB model, perfused it with control or *Pf-*PBMC, and performed multimodal analyses, including scRNA-seq, FACS, and Luminex multiplex assay (Figure 2A and S2A). Briefly, the 3D-BBB model is composed of a collagen hydrogel containing primary brain astrocytes and pericytes, crossed by a microfluidic network, and then seeded with endothelial cells that generate a 3D tube-like structure. The devices are composed by a 13 by 13 network that presents a wide gradient of flow conditions in a single device, with high flow velocity and wall shear stress near the inlet and the outlet, and low velocity and sheer stress in the opposite corners.^15,26,18^ After fabrication, the 3D-BBB model was cultured for 6 days, during which cells reach a peak in BBB marker expression and barrier function, a significant improvement from endothelial-only 3D models.^18^ After stimulation, control and *Pf-*PBMC were resuspended in fresh media, removing all soluble iRBC remnants, at 3 x 10^6^ PBMC/ml, which approximates physiological immune cell blood concentrations.^27^ Control or *Pf-*PBMC were perfused for 1-hour under gravity-driven flow, and 3D-BBB devices were incubated with the cytoadherent leukocytes for 10 hours before processing for downstream analysis.

**Figure 2.**
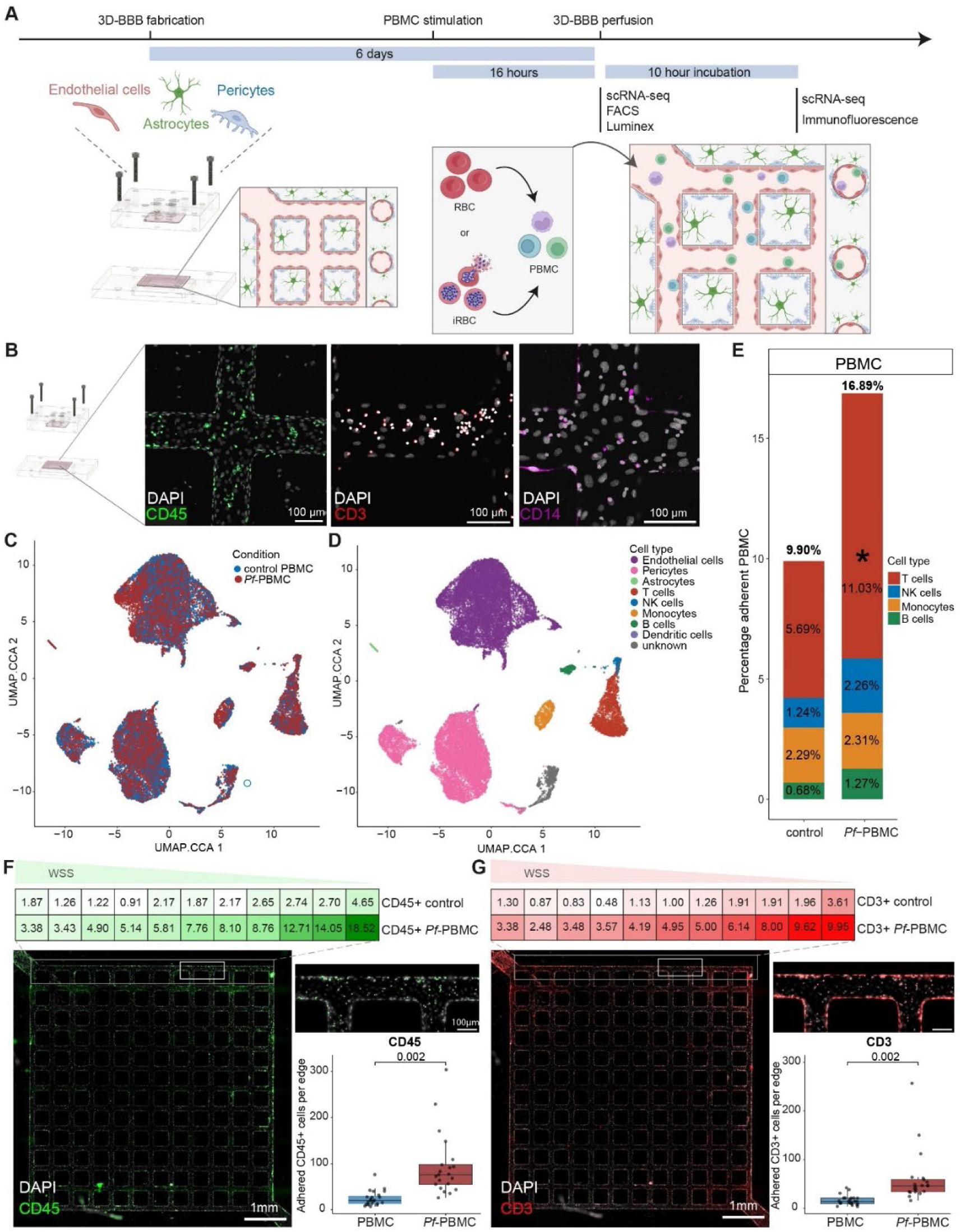
*P. falciparum*-stimulated PBMC present increased binding to the 3D-BBB model, specifically in *P. falciparum*-stimulated T cells. **A.** Schematic overview of the experimental workflow. **B.** Representative confocal imaging (maximum z-projection) of perfused 3D-BBB devices showing adherent leukocytes stained for CD45 (green), T cells stained for CD3 (red), and monocytes stained for CD14 (pink). Nuclei are labelled with DAPI (white). **C-D.** UMAP visualization of single-cell transcriptomic data from perfused 3D-BBB devices, colored by experimental condition (**C**) and annotated cell type identity (**D**). **E**. Bar plot showing the proportion of each immune cell type adherent to the 3D-BBB model in control and *Pf*-stimulated conditions. Asterisks labels statistically significant differences identified by *scCODA* compositional analysis (FDR < 0.05). **F-G**. Quantification and representative images of adherent leukocytes stained for CD45 (green) (**F**) and T cells stained for CD3 (red) (**G**). Each point represents one edge quantification coming from n=3 independent PBMC donors (two 3D-BBB devices per donor per condition), Mann Whitney U test. Heatmaps show number of adherent cells per quantification region of interest (ROI) (see Figure S2E).

Cytoadherence of the perfused PBMC was confirmed by confocal immunofluorescence microscopy and staining for the pan-leukocyte marker CD45, the T cell marker CD3, and the monocyte marker CD14 in the 3D-BBB model (Figure 2B). To better characterize the populations of leukocytes that interact with the microvessels, the 3D-BBB devices were dissociated for subsequent scRNA-seq analysis. This approach captures both BBB cells and the cytoadherent PBMC population, and excludes the leukocytes that did not bind. After quality control, we obtained data from 29,010 cells derived from 3D-BBB devices perfused with control and *Pf-*PBMC from 3 donors. Using a UMAP, we visualized the clustering of different cell types present in the 3D model, identified by unsupervised clustering and annotated using known cell type marker genes (Figures 2C, 2D, S2B, S2C). The two largest populations represented endothelial cells and pericytes, while only a very small population of astrocytes could be confidently identified. A fourth population of BBB cells was classified as unknown due to its mixed expression of astrocyte, fibroblast, and glial cell markers. Notably, we identified a considerable number of PBMC that bound to the 3D-BBB model in both conditions, with T cells representing the largest population (Figure 2D).

Previous studies suggest that immune cell accumulation in the microvasculature of CM patients is primarily driven by inflammatory upregulation of endothelial receptors like ICAM-1 and VCAM-1.^13,28^ Notably, even without pre-stimulation of the 3D-BBB models with inflammatory cytokines, the scRNA-seq data revealed almost a doubling in binding of *Pf*-PBMC compared to control PBMC, with adherent leukocytes making up 16.87% or 9.90% of all sequenced cells, respectively (Figure 2E). *P. falciparum*-stimulated T cells exhibited a significant increase in binding from 5.69% to 11.03%, identified using the *scCODA* Bayesian model for compositional single-cell data analysis.^29^ Stimulated B cells and NK cells showed non-significant increases, whereas monocyte accumulation was unchanged (Figures 2E and S2D).

The scRNA-seq analysis revealed a near doubling in binding of *Pf-*PBMC relative to the total cell number in 3D-BBB model, mostly consisting of vascular cells. To determine the absolute increase in cytoadhesion, we imaged CD45+ (pan-leukocyte marker) and CD3+ (T cell marker) adherent leukocytes across the gradient of flow states present in the 3D-BBB model. An automated quantification pipeline was used to threshold, overlay, and count individual cell immunofluorescence signals along the outer edges of the 3D-BBB model (Figure S2E). The binding of both CD45+ and CD3+ cells was significantly increased, with leukocytes showing a 3.8-fold increase (20 and 76 cells per edge; Interquartile range (IQR), 13-30 and 55-98 for control and *Pf*-PBMC, respectively) and T cells demonstrating a similar 3.3-fold increase (14 and 46 cells per edge; IQR, 10-21 and 34-59 for control and *Pf*-PBMC, respectively) (Figure 2F and G). Interestingly, this binding increase was wall shear stress-dependent as leukocytes demonstrated a 2.7-fold binding increase in high-shear stress regions, and a 4.5-fold binding increase in low-shear stress regions (Figure 2F and G). These results were consistent throughout three different PBMC donors (Figures S2F and S2G). Taken together, our results identified that *P. falciparum*-stimulation of leukocytes increases their adherence to endothelial cells, an effect particularly marked in T cells.

### *P. falciparum-*stimulated CD8+ effector memory and γδ T cells present increased binding to the 3D-BBB model

Given that T cells encompass various subpopulations with putative roles in CM pathogenesis, including CD8+ T cells,^9,11,30–32^ CD4+ T cells,^33,34^ and γδ T cells,^35–37^ we performed a more detailed characterization of the T cell subsets enriched in the 3D-BBB model after *P. falciparum-*stimulation. By mapping the adherent PBMC (Figure 2D) to the unperfused PBMC scRNA-seq dataset reference (Figure 1C), we obtained a high-confidence annotation of adherent T cell subpopulations despite their low numbers. Although all *P. falciparum*-stimulated T cell subsets showed increased binding, only CD8+ TEMs and γδ T cells exhibited significant ∼3-fold increases (Figure 3A, S3A). To determine whether enhanced binding could be a consequence of T cell activation, we focused on the state of *Pf-*PBMC before perfusion. We first defined a *T cell cytotoxic and effector* signature consisting of general cytotoxic markers (*GZMB, PRF1, CD69, TNFRSF9, LAG3, TNFRSF4, CRTAM, NKG7*), and effector cytokines (*TNF, IFNG*) known to be upregulated upon *P. falciparum-*activation and to contribute to CM pathogenesis.^9,24^ Indeed, both CD8+ TEMs and γδ T cells displayed the strongest increase in activation marker expression after exposure to *P. falciparum-*iRBC (Figure 3B). While the observed binding enhancement could be attributed to changes on either endothelial cells or T cells, endothelial cells remained in a resting state without increased surface expression of the ICAM-1 endothelial receptor during the 1-hour perfusion timeframe (Figure S3B). Therefore, we propose that *P. falciparum*–induced alterations in T cells promote their enhanced endothelial binding.

**Figure 3.**
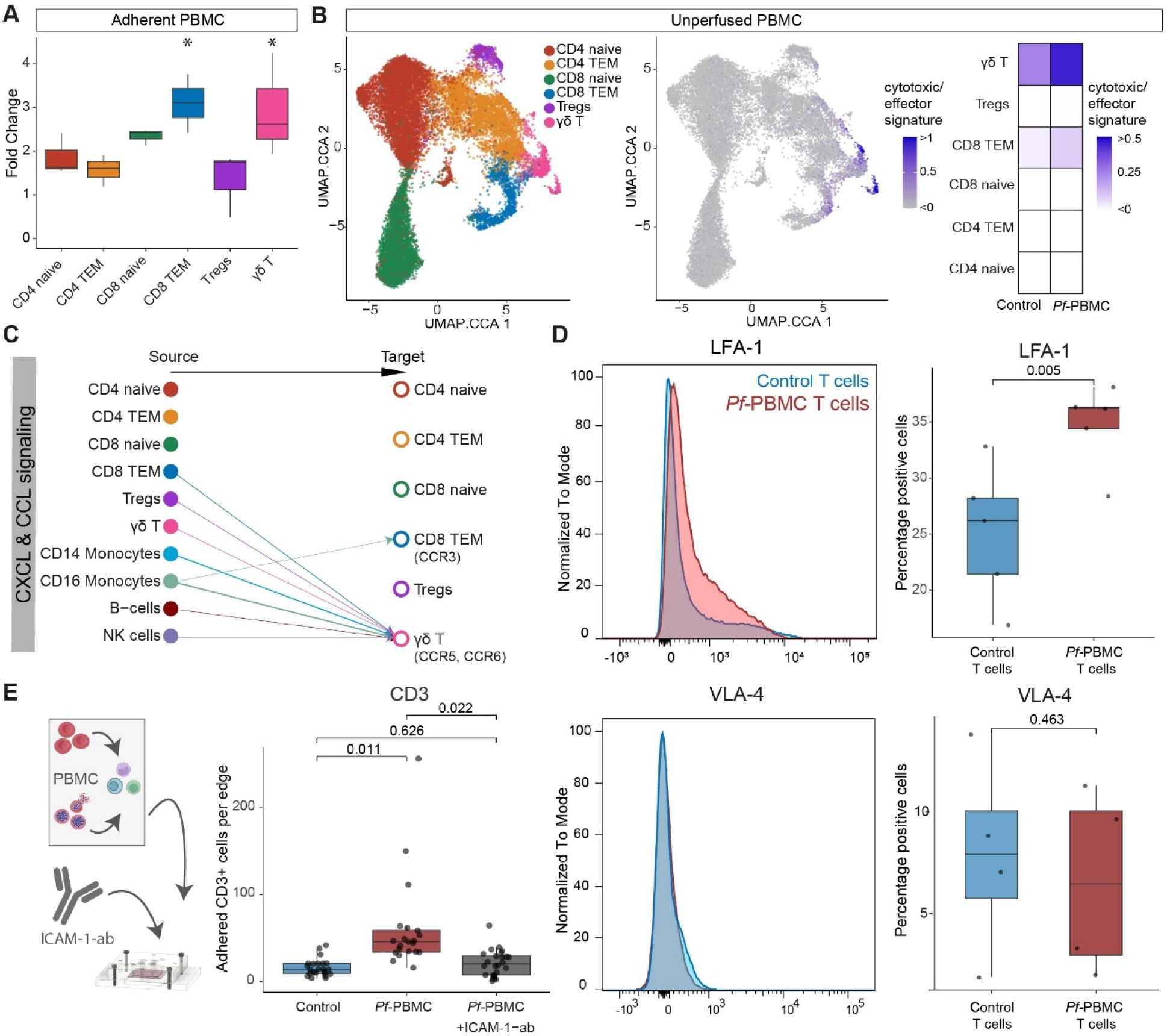
*P. falciparum*-activated T cell subpopulations present increased endothelial binding mediated by a switch to high-affinity LFA-1 integrin conformations. **A.** Fold change in the abundance of adherent T cell subpopulations between *P. falciparum*-stimulated and control conditions. Asterisks label statistically significant differences identified by *scCODA* compositional analysis (FDR < 0.15). **B.** UMAP visualization of T cells, colored by annotated subpopulation (left) and by *cytotoxicity/effector gene signature scores* (middle) and heatmap comparing *cytotoxicity/effector gene signature scores* across T cell subpopulations in control and *Pf-*PBMC conditions (right). **C.** Chemokine receptor–ligand interactions (CXCL, CCL pathways) towards T cells in *Pf*-PBMC, identified using the *CellChat* package. Arrows point from sender cells to receiver cells and are colored by sender cell. Arrow thickness is proportional to the interaction strength. **D.** Representative flow cytometry histograms showing expression levels of high binding affinity conformations of LFA-1 (top) and VLA-4 (bottom) in T cells from *P. falciparum*-stimulated (red) and control (blue) conditions and quantification of percentage of LFA-1 (top) or VLA-4 (bottom) -positive T cells. Histograms display fluorescence intensity (x-axis) and relative cell count (y-axis), normalized to mode for distribution comparison. Each dot represents an independent FACS experiment with a different PBMC donor (n = 4-5). Paired Student’s t-test. **E.** Number of adherent CD3+ T cells per edge of the 3D-BBB model with a 30-minute pre-treatment of the devices with a monoclonal ICAM-1-blocking antibody, compared to untreated devices perfused with control and *Pf*-PBMC T cells (same data as Fig. 2G). Each point represents one edge quantification coming from n=3 independent PBMC donors (two 3D-BBB devices per donor per condition), Kruskal– Wallis test with Dunn’s pairwise comparisons test and BH correction.

T cell activation can lead to changes in conformation and affinity of various leukocyte integrins, including LFA-1 and VLA-4,^38–40^ which enhance leukocyte binding to endothelial cells. Conformational changes in leukocyte integrins can be induced by T cell receptor (TCR) or chemokine signaling.^38,39,41^ Indeed, *CellChat* interaction analysis on the pre-perfusion *Pf-*PBMC dataset revealed that chemokine signaling was largely directed towards γδ T cells, followed by limited signaling towards CD8+ TEM and no evident signaling to other T cell subsets (Figures 3C and S3C). We assessed the level of the conformationally active high-affinity forms of the VLA-4 and LFA-1 integrins by FACS analysis. While we could not observe any changes in VLA-4, there was a significant shift towards the high-affinity LFA-1 conformation on the surface of T cells after 16-hour stimulation with *P. falciparum-*iRBC (Figures 3D). To assess whether LFA-1 contributes to increased T cell binding to endothelial cells, we inhibited the interaction with its cognate endothelial receptor through pre-incubation of the 3D-BBB model with a monoclonal ICAM-1 blocking antibody. This resulted in a significant reduction in T cell binding, nearly to levels observed in control T cells (Figure 3E and S3D), highlighting the role of LFA-1 – ICAM-1 interactions. Taken together, these findings suggest that the accumulation of *P. falciparum*-stimulated T cells in the 3D-BBB model is primarily driven by activated CD8+ TEM and γδ T cells, and is associated with LFA-1 conformational changes, potentially induced by chemokine signaling from other leukocyte subsets.

### *P. falciparum-*stimulated leukocytes induce inflammatory activation of BBB cells

Disentangling the contribution of host immune response or parasite to microvascular injury remains difficult in patients and animal models. In contrast, our modular bioengineered system enables selective analysis of leukocyte effects on the BBB, independent of iRBC sequestration. To identify potential immune-driven disruptive mechanisms, we profiled transcriptomic and functional responses of the BBB cells after exposure to *Pf-*PBMC. Weighted gene co-expression network analysis (WGCNA) identifies correlated gene expression patterns in high-dimensional data, grouping genes into transcriptional modules that represent coordinated biological processes.^42–44^ We applied this analysis as an unbiased method to identify shared altered processes across BBB cell types as well as pathway alterations specific to endothelial cells. Of the 6 identified transcriptional modules in the 3D-BBB immunocompetent model, three were found to be differentially expressed between the *Pf*-PBMC and the control condition (module 1, 5, and 6; expression in > 30% of cells, log2FC > 2) (Figures S4A and S4B). Module 6 was shared and differentially expressed in both brain endothelial cells and pericytes, while the two other modules were specific to endothelial cells (Figures 4A and S4B). GO-term over-representation analysis on the top hub genes in module 6 revealed involvement in innate inflammatory immune responses and defense mechanisms against foreign organisms, suggesting an upregulation of a shared inflammatory gene program in BBB cells (Figure 4B). Concordantly, we found similar results in conventional differential gene expression analysis. Endothelial cells and pericytes presented an overlap in DE genes (Figure 4C, S4C), and GO-term over-representation highlighted upregulation of immune-related processes, including cytokine and chemokine signaling, interferon response, antigen presentation, NF-κB and JAK/STAT signaling (Figure 4D). The inflammatory activation in the 3D-BBB model was confirmed through ICAM-1 immunofluorescence staining, showing a significantly increased expression in brain endothelial cells, pericytes, and astrocytes after the 10-hour *Pf-*PBMC incubation (Figure 4E). High-content imaging in 2D monolayers confirmed significant ICAM-1 induction by *Pf-*PBMC from six different donors, comparable to TNF-α at 10ng/ml (Figure S4D). Notably, exposure to matched doses of *P. falciparum-*iRBC derived products did not upregulate ICAM-1, indicating that the *Pf*-PBMC effect is not attributable to residual iRBC products (Figure S4D). Overall, two orthogonal analysis methods, unsupervised WGCNA and differential gene expression analysis uncovered a shared inflammatory signature induced by *Pf*-PBMC in BBB cell, which could be validated using immunofluorescence.

**Figure 4.**
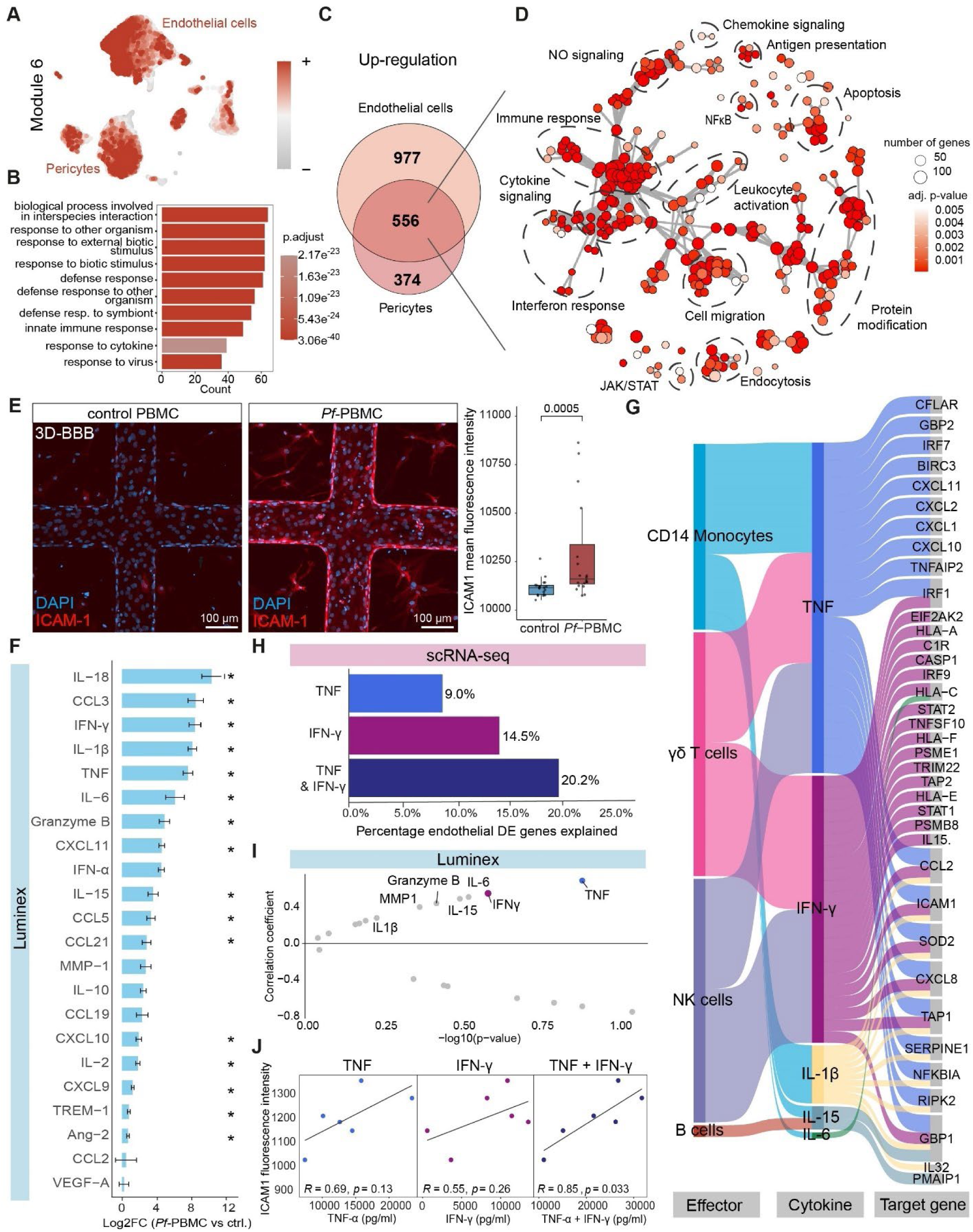
Monocytes, γδ T cells, and NK cells induce inflammatory activation of BBB cells, mainly driven by TNF-α and IFN-γ. **A-B**. Transcriptional module 6, identified *via* weighted gene co-expression network analysis (WGCNA), is differentially expressed in endothelial cells and pericytes after *P. falciparum*-PBMC exposure (log2FC > 2 and expression in >30% of cells) (Figure S4A, B). UMAP plot shows module 6 eigengene expression (**A**). Gene Ontology (GO) term over-representation analysis on the top 100 hub genes (ranked by eigengene-based connectivity, kME), ordered by gene count and colored by adjusted p-value (**B**). All transcriptional modules are shown in Figures S4A, B. **C.** Venn diagram of significantly upregulated genes in endothelial cells and pericytes (FDR < 0.05, log2FC > 0.05) (Figure S4C). **D.** GO term over-representation analysis of 556 overlapping upregulated DE genes in endothelial cells and pericytes. Network nodes represent the most significant GO-terms and edges connect GO-terms with more than 30% gene overlap. GO-term clusters were manually labelled. **E.** Representative confocal maximum intensity z-projection showing ICAM-1 (red) labeling in 3D-BBB models after perfusion and 10-hour incubation with control (left) and *Pf*-PBMC (middle) and quantification of mean pixel fluorescence intensity (right). Each point represents one ROI coming from n=4 independent PBMC donors (one 3D-BBB device per donor per condition), Mann Whitney U test. **F.** Log2FC increase in cytokine levels measured by Luminex in supernatants of control or *Pf-*PBMC. Asterisks denote significance with paired Student’s t-test and Benjamini–Hochberg (BH) correction (n = 6 per condition). **G.** Integration of Luminex cytokine data with scRNA-seq data using *Nichenet* ligand-target matrix. Flow diagram shows cytokine–target gene relationships between immune cell types (cytokine producers) and endothelial cells (target gene responders). Links are based on *Nichenet* regulatory potential scores (> 0.2) between significantly increased cytokines (Luminex) and significantly upregulated genes in endothelial cells (scRNA-seq, FDR < 0.05, log2FC > 0.05). See details in Methods. **H.** Proportion of significantly upregulated genes in endothelial cells (FDR < 0.05, log2FC > 0.05) predicted to be regulated by TNF-α or IFN-γ. Regulation was inferred if TNF-α or IFN-γ were among the top 5 predicted regulators of a given gene in the *Nichenet* matrix. **I.** Correlation analysis between absolute cytokine levels (Luminex) in the supernatant of *Pf*-PBMC from n = 6 PBMC donors and induced ICAM-1 expression levels in endothelial monolayers. The x-axis shows – log_10_(p-value) and the y-axis shows Pearson correlation coefficients. Cytokines with the strongest positive correlations are labeled. **J.** Scatter plots showing the correlation between TNF-α and IFN-γ levels (individually and summed) from n = 6 PBMC donors on the x-axis and induced ICAM-1 fluorescence intensity in endothelial cells on the y-axis. Pearson correlation coefficient (R) and p-values are indicated.

### Increased secretion of TNF-α and IFN-γ by monocytes, γδ T cells, and NK cells induces BBB activation

Increased levels of pro-inflammatory cytokines is a common feature in blood and brain samples of CM patients.^24,45–48^ We therefore aimed to identify the cell types and effector proteins responsible for the broad inflammatory activation phenotype affecting BBB cells in the 3D model. Luminex profiling of supernatants from control and *Pf-*PBMC co-cultures from 6 different donors showed significant increases in interleukin (IL)-18, CCL3, IFN-γ, IL-1β, TNF-α, IL-6, and granzyme B (Figure 4F), among other cytokines known to be elevated in the plasma of CM patients.^24,45,49–52^ To discern which of the upregulated cytokines most strongly activates BBB cells, we integrated the Luminex and scRNA-seq datasets using *Nichenet*,^53^ filtering for cytokine ligands increased by Luminex and significantly upregulated endothelial target genes. The integration revealed TNF-α and IFN-γ as the dominant regulators of the induced endothelial target genes, while IL-1β, IL-15, and IL-6 only contributed to few (Figure 4G). Remarkably, 20.2% of the entirety of significantly upregulated endothelial genes could be attributed to the signaling response of just two cytokines, TNF-α and IFN-γ (Figure 4H). Conversely, all the remaining increased cytokines combined were only responsible for 21.3% of significantly upregulated genes. Using the scRNA-seq dataset, we identified CD14+ monocytes, γδ T cells, and NK cells as the primary effector cell types secreting both cytokines (Figure 4G). Correlation between the full cytokine panel and ICAM-1 fluorescence intensities in 2D endothelial monolayers revealed TNF-α and IFN-γ as the strongest inflammatory correlates (TNF-α: R = 0.69 / p = 0.13, IFN-γ: R = 0.55 / p = 0.26; Figures 4I, 4J and S4E), with the association becoming significant when the concentrations of both cytokines were added up (R = 0.85 / p = 0.033) (Figure 4J). Consistently, a linear regression model including both TNF-α and IFN-γ significantly improved the fit for ICAM-1 levels compared with either cytokine alone (likelihood ratio tests: p = 0.053 and p = 0.019, compared to TNF-α or IFN-γ alone respectively). Similar strong correlations between TNF-α and IFN-γ expression and inflammatory DE genes were found in the scRNA-seq analysis of the 3D-BBB model (Figure S4F). Taken together, the BBB cell types exhibited robust inflammatory activation driven by TNF-α and IFN-γ secretion from *P. falciparum*-stimulated CD14 monocytes, γδ T cells, and NK cells.

### *P. falciparum*-stimulated PBMC induce endothelial cytoskeletal changes and apoptosis, resulting in BBB disruption

Although reports support the accumulation of CD8+ T cells^9,10^ and monocytes^8^ in the brain microvasculature of CM, their role in BBB damage during CM remains largely unknown. WGCNA identified two modules (module 1 and 5) differentially expressed exclusively in endothelial cells upon *Pf-*PBMC exposure (Figures S4A and S4B). GO-term over-representation analysis on the module top hub genes revealed processes suggestive of endothelial barrier dysfunction, including cytoskeletal, phosphorylation, and cell motility processes in module 1, and apoptotic processes in module 5 (Figure 5A). Staining for the endothelial actin cytoskeleton revealed a significant induction of stress fibers, indicative of a contractile and barrier destabilizing phenotype (Figure 5B). These cytoskeletal changes were accompanied by loss of the junctional marker VE-cadherin (Figures 5C and S5A), and elevated cleaved caspase 3 levels, suggestive of apoptosis (Figure 5D). We next tested whether these changes result in functional barrier disruption by measuring permeability to 70 kDa fluorescent dextran in the 3D-BBB model and calculating the ratio of values obtained 10 hours post-PBMC-incubation relative to the pre-perfusion baseline. *Pf-*PBMC from three independent donors significantly increased the permeability ratio in the 3D-BBB model, demonstrating that the transcriptional changes translate into functional BBB disruption (Figure 5E). Consistently, measurements on 2D endothelial monolayers by real-time impedance assays (xCELLigence) using six additional donors support a robust decline in endothelial barrier integrity in response to *Pf-*PBMC (Figures S5B and S5C). Control PBMC did not significantly affect barrier integrity in either 2D or 3D settings, indicating that the disruptive phenotype is not driven by allogenic responses between PBMC and endothelial cells (Figures 5E and S5C). Direct exposure to *P. falciparum* iRBC–derived products at concentrations equivalent to the PBMC stimulation did not change 2D monolayer integrity, confirming that residual parasite products are not the cause for barrier disruption (Figure S5C). Altogether, exposure to *Pf*-PBMC triggers endothelial cytoskeletal and apoptotic pathways which results in BBB disruption.

**Figure 5.**
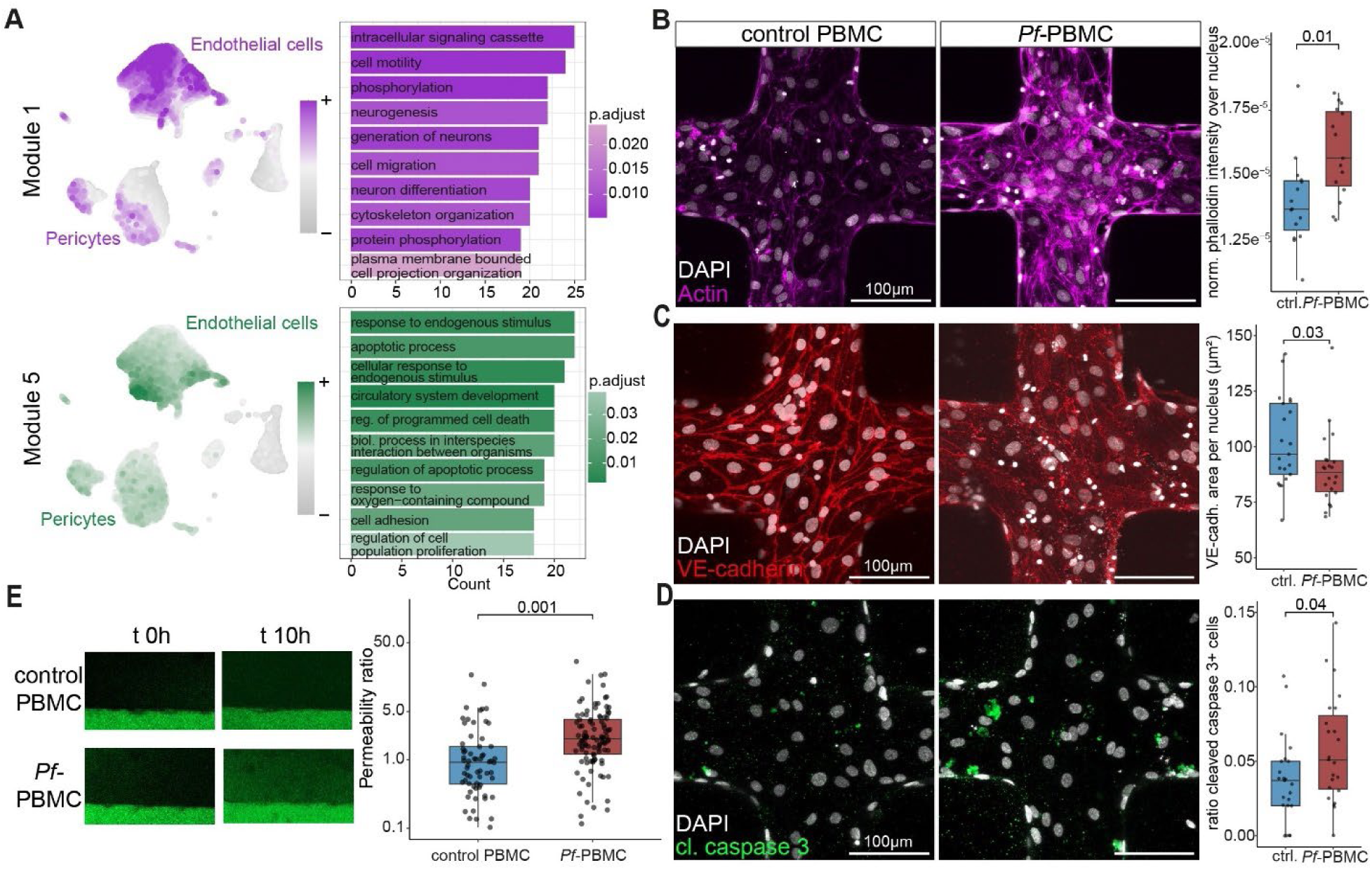
*P. falciparum*-stimulated PBMC induce upregulation of cytoskeletal and apoptotic transcriptional modules, and disrupt BBB integrity in the 3D-BBB model. **A.** UMAP visualization of eigengene expression for transcriptional modules 1 and 5, identified by WGCNA as significantly upregulated in endothelial cells exposed to *Pf*-PBMC versus control (left; log2FC > 2 and expression in > 30% of cells) and GO-term over-representation analysis on the top 100 hub genes (right; ranked by eigengene-based connectivity, kME). All identified modules can be found in Figures S4A, B. **B.** Representative confocal maximum intensity z-projection showing phalloidin (magenta) staining of F-actin in 3D-endothelial monoculture models after perfusion and 10-hour incubation with control or *Pf*-PBMC. Quantification of pixel mean fluorescence intensity over the nucleus normalized to total cellular intensity (right). **C.** Representative confocal maximum intensity z-projection showing VE-cadherin (red) staining in 3D-BBB devices after perfusion and 10-hour incubation with control or *Pf*-PBMC. Quantification of segmented junctional VE-cadherin area per endothelial cell (right) (see also Figure S5A). **D.** Representative confocal maximum intensity z-projection showing cleaved caspase-3 (green) labeling in 3D-BBB devices after perfusion and 10-hour incubation with control or *Pf*-PBMC. Quantification of proportion of cleaved caspase-3-positive endothelial cells (right). **B.-D.** Each point represents one ROI coming from n=3 independent PBMC donors (one 3D-BBB device per donor per condition), Mann Whitney U test. **E.** Representative images and quantification of 70 kDa FITC-dextran perfusion in 3D-BBB devices. Apparent permeability ratios were calculated by comparing 10-hour post-incubation to baseline (pre-incubation) measurements. Each point represents an ROI from 3D-BBB models exposed to *Pf*-PBMC (n = 7 independent devices, 3 PBMC donors) or control PBMC (n = 6 independent devices, 3 PBMC donors).

### BBB disruption caused by *P. falciparum*-stimulated PBMC is dependent on direct endothelial-leukocyte binding

TNF-α and IFN-γ emerged as key drivers for inflammatory BBB cell activation in the 3D model. However, the presence of granzyme B expressing CD8+ T cells in CM brain postmortem samples^9^ argues for the existence of additional cytotoxic effector proteins that could mediate barrier disruption. We used the Luminex data on PBMC-secreted proteins to evaluate their disruptive potential. Notably, granzyme B and IFN-γ presented the strongest correlations with 2D xCELLigence barrier measurements across 6 independent donors (r = 0.88 and r = 0.81, respectively; Figure 6A and S5D), while other cytokines, such as TNF-α that showed a prominent role in BBB inflammation, were not associated with barrier disruption. Analysis on the scRNA-seq dataset revealed significantly increased gene expression of granzyme B and IFN-γ in *P. falciparum*-stimulated γδ T cells and NK cells (Figures 6B and C). To unravel locally confined from global disruptive effects mediated by *Pf-*PBMC-secreted proteins, we subclustered the endothelial cells from the *Pf*-PBMC condition and inferred ligand-receptor communications between endothelial cells and *Pf*-PBMC using *CellChat* (Figure 6D and S5E). The analysis identified three endothelial subclusters (clusters 2, 10, and 12) showing the highest predicted interaction probabilities with leukocytes (Figure 6D). To evaluate the degree of disruption across endothelial clusters, we computed a *combined disruptive module score* including the top hub genes from transcriptional modules 1, 5, and 6 (Figures 4A, 5A, S4A and S4B). *Disruptive module score* values peaked in endothelial clusters 2, 10 and 12, corresponding to those with the strongest predicted interactions with *Pf-*PBMC (Figure 6E and S5F). This result indicates that *Pf-*PBMC sequestration hotspots might coincide with maximal transcriptional disruption.

**Figure 6.**
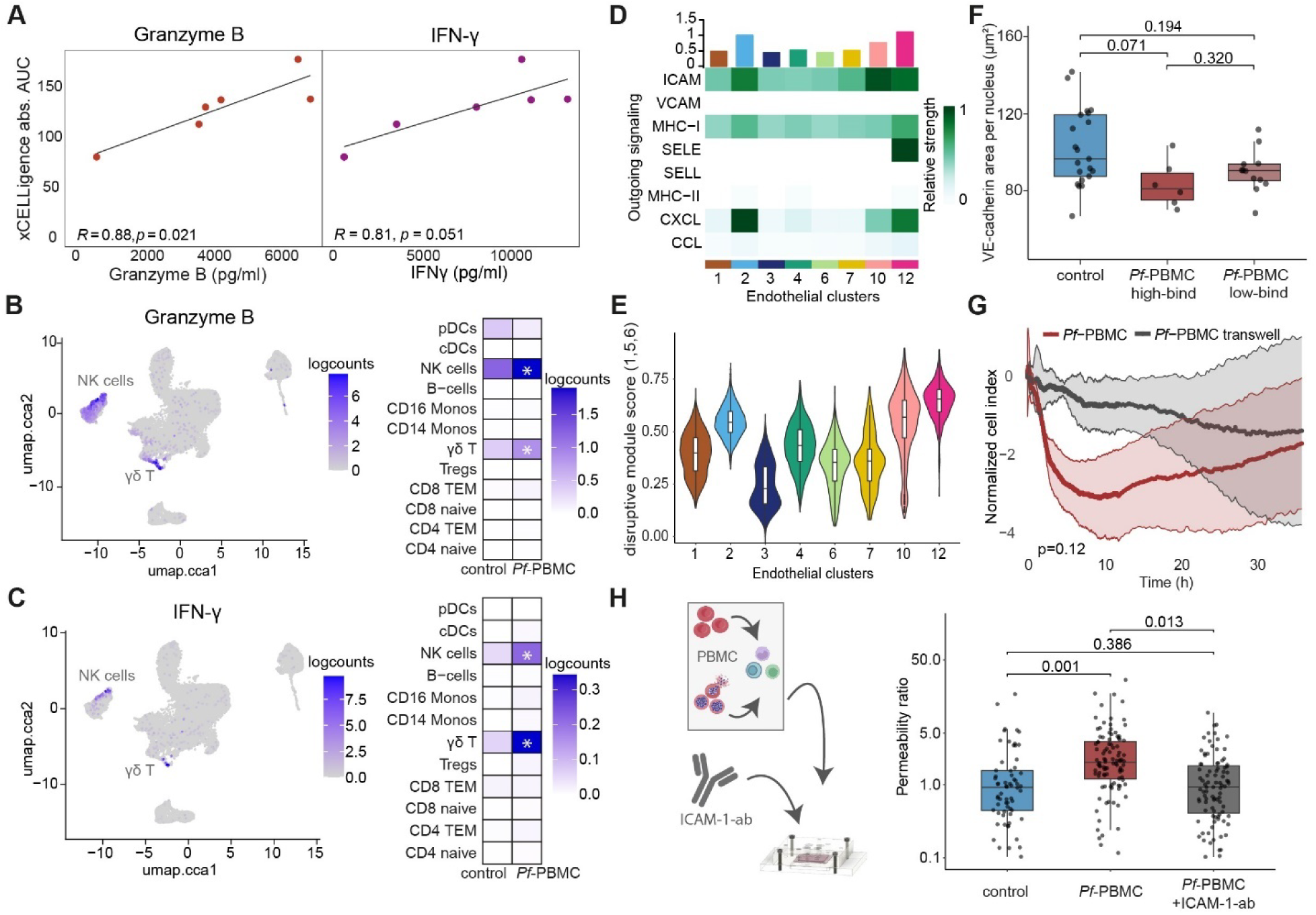
BBB disruption induced by *P. falciparum*-stimulated PBMC is dependent on immune cell binding to the endothelium. **A.** Scatter plots showing the correlation between granzyme B and IFN-γ levels (x-axis; Luminex data from n = 6 PBMC donors) and endothelial barrier disruption (y-axis; area under the curve (AUC) from xCELLigence impedance measurements). Pearson correlation coefficients (R) and p-values are shown. **B-C.** UMAP visualization and heatmap showing the granzyme B (**B**) and IFN-γ (**C**) expression levels in PBMC between *P. falciparum*-stimulated and control condition **D.** Heatmap showing relative strength of selected signaling pathways (ICAM, VCAM, MHC-I, MHC-II, SELE (E-selectin), SELL (L-selectin), CXCL, CCL) between *Pf*-PBMC and endothelial cells stratified by endothelial cluster as identified by *CellChat* analysis. **E.** Violin plots showing the combined expression scores of transcriptional modules 1, 5, and 6 (top 100 hub genes ranked by eigengene-based connectivity, kME) in *Pf*-PBMC-exposed endothelial cells across subclusters identified by unsupervised clustering (see Figure S5E). **F.** Quantification of VE-cadherin junctional area in regions of high-versus low-*Pf-*PBMC binding within the 3D-BBB model. Each point represents one ROI coming from n=3 independent PBMC donors (one 3D-BBB device per donor per condition), Kruskal–Wallis test with Dunn’s pairwise comparisons test and BH correction. **G.** Real-time trans-endothelial electrical impedance measurements using xCELLigence to assess barrier integrity in endothelial monolayers following exposure to *Pf*-PBMC, with or without direct cell contact (transwell insert). Cell index is normalized to the time point immediately before PBMC addition and the respective control PBMC condition. n=3 independent PBMC donors and experiments, Paired Student’s t-test. **H.** Apparent 70 kDa FITC-dextran permeability ratio (10-hour post-incubation vs. baseline) in 3D-BBB models exposed to control or *Pf*-PBMC (same data as Fig. 5E), or *Pf-*PBMCs following 30-minute pre-treatment of the 3D-BBB with a monoclonal ICAM-1–blocking antibody. Each point represents an ROI from 3D-BBB models exposed to *Pf*-PBMC (n = 7 independent devices), control PBMC (n = 6 independent devices), or *Pf*-PBMC after monoclonal ICAM-1-blocking antibody perfusion (n = 6 independent devices) (two 3D-BBB devices per donor per condition). Kruskal–Wallis test with Dunn’s pairwise comparisons test and BH correction.

To further disentangle soluble from contact-dependent effects of secreted effector proteins, we leveraged the microfluidic design of the model that creates high- and low-*Pf-*PBMC binding regions (Figures 2F and G). We stratified the VE-cadherin junctional area in the 3D-BBB model by low-versus high-PBMC binding regions. Regions of high *Pf-*PBMC binding showed a greater loss of VE-cadherin, although not reaching significance (Figure 6F). Further evidence for *Pf*-PBMC binding-dependent disruption included xCELLigence barrier measurements, showing minor disruption when *Pf-*PBMC were placed in a transwell without physical contact, compared to a rapid decline in endothelial barrier integrity when leukocytes were in direct physical contact with endothelial cells (p-value = 0.12) (Figure 6G). Given accumulated evidence that *Pf*-PBMC-induced BBB disruption is contact-dependent, we performed functional validation in the 3D-BBB model. We pre-incubated the 3D-BBB model with an ICAM-1-blocking monoclonal antibody before *Pf*-PBMC perfusion and quantified 70 kDa permeability before and after the 10-hour incubation period. ICAM-1 blockade reduced *Pf*-PBMC binding to control PBMC levels (Figure 5E and S5G), and completely abrogated the permeability increase in the 3D-BBB model (Figure 6H), indicating that enhanced *Pf*-PBMC adhesion and direct endothelial contact are required for BBB disruption.

### *P. falciparum* products and *P. falciparum*-stimulated PBMC induce shared and distinct endothelial disruptive pathways of CM pathogenesis

Our data illustrate the importance of *Pf-*PBMC-driven BBB pathology. Nevertheless, the sequestration of *P. falciparum-*iRBC remains a defining hallmark of disease.^7^ We therefore compared endothelial pathogenic programs elicited by these two key drivers, by integrating our current scRNA-seq data with data generated in the same 3D-BBB model after exposure to iRBC-derived products.^18^ This analysis enables to identify major pathogenic molecular mechanisms, attributes the resulting phenotypes to parasite- or leukocyte-driven mechanisms, and infers the effector cell type within the model (Figure 7A). We focused on endothelial transcriptomic changes, given the role of this cell type as the main interface with parasites and circulating leukocytes. Integrated GO-term over-representation analysis revealed shared dysregulated processes across both stimuli, as well as stimulus-specific responses unique to parasite- or leukocyte-exposure (Figure 7B). The shared processes could represent key pathways in CM pathogenesis, offering valuable insights for the development of CM targeted treatment strategies. These included NF-κB-signaling (e.g. *NFKBIA, CD40, MYD88*), antigen-presentation (e.g. *HLA-A/B/C/E/F*), interferon-induced genes (e.g. *IFIT1, ISG15, GBP1/2*), and the JAK-STAT signaling pathway (e.g. *STAT1/2/3/6, JAK2*). Overall, JAK-STAT emerges as a dominant dysregulated pathway, driven by both parasite and leukocytes, and upregulated across all effector cells, including BBB cells and all immune cell types (Figure 7C). Another commonly activated pathway is NF-κB-signaling, indicated by ICAM-1 induction in endothelial cells and pericytes, following both iRBC and *Pf-*PBMC exposure (Figure 7D).

**Figure 7.**
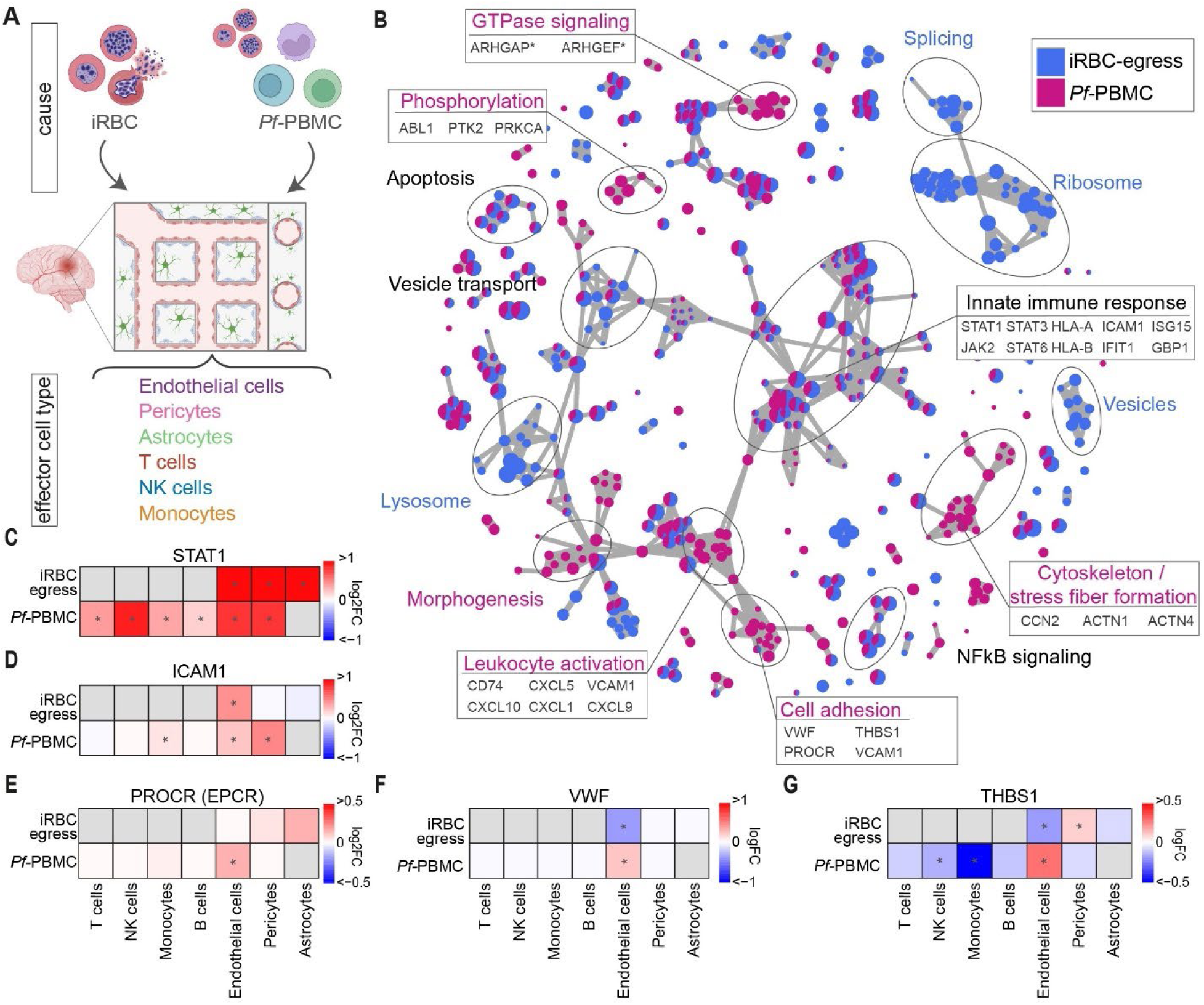
*P. falciparum-*iRBC and immune cell induce shared and distinct disruptive pathways in the 3D-BBB model. **A**. Schematic overview of the datasets included in the integrative analysis, namely perfusion of the 3D-BBB model with iRBC-egress products (compared to uninfected RBC control) and *Pf-*PBMC (compared to control PBMC). **B**. Comparative GO-term over-representation analysis on upregulated genes in endothelial cells (FDR < 0.05, log2FC > 0.05) upon exposure to *Pf-*PBMC or iRBC-egress products. Selected processes and genes only upregulated upon exposure to *Pf-*PBMC (magenta), *P. falciparum-*iRBC (blue), or shared (black font). **C-G**. Heatmaps showing *STAT1* (**C**), *ICAM1* (**D**), *PROCR* (EPCR) (**E**), *VWF* (**F**), and *THBS1* (**G**) log2FC changes in endothelial cells, pericytes, astrocytes, T cells, NK cells, monocytes, and B cells in the *Pf-*PBMC or the iRBC-egress datasets compared to the respective controls. Asterisks labels statistically significantly differentially expressed genes (FDR < 0.05, log2FC > 0.1).

By contrast, the analysis also identified genes and pathways that were specifically induced by either *Pf-*PBMC or iRBC. While iRBC-egress distinctly induced an increase in splicing, ribosomal, lysosomal, and vesicle related processes,^18^ *Pf*-PBMC induced alterations in leukocyte activation, cytoskeleton, and cell adhesion related processes as well as phosphorylation, and GTPase signaling (Figure 7B). Among the *Pf-*PBMC-specific upregulated endothelial responses, we identified genes known to be involved in leukocyte activation and recruitment during CM, including *CXCL9* and *CXCL10*, which are strongly upregulated by IFN-γ.^54^ We also identified *Pf-*PBMC-induced upregulation of several genes mediating endothelial interactions with *P. falciparum-*iRBC either directly or indirectly, such as *PROCR, VCAM-1, THBS1,* and *VWF*. Of special relevance is the EPCR-encoding gene *PROCR*, a receptor implicated in endothelial sequestration of iRBC^55^ and highly associated with CM severity^56–58^ (Figure 7E), found to be selectively upregulated by *Pf-*PBMC in endothelial cells. Likewise, vWF, a central component of the coagulation cascade that contributes to platelet clot formation^59^ and has been implicated in indirect binding to *P. falciparum-*iRBC through platelets,^60^ was also specifically induced by *Pf-*PBMC (Figure 7F). Similarly, Thrombospondin (*THBS1*) showed *Pf-*PBMC driven, endothelial-specific upregulation (Figure 7G). This protein was initially suggested to act as an iRBC receptor^61,62^ and a THBS1 haplotype has been linked to malaria susceptibility.^63^ As a proof-of-concept on how to link our current analysis with population-based association studies, we included predictions from the cell type-specific enhancer-gene prediction model ENCODE-rE2G.^64^ The model could map an intronic single nucleotide polymorphism (SNP) from the malaria-associated THBS1 haplotype to an enhancer region predicted to be selectively active in brain microvascular endothelial cells. In combination with our scRNA-seq results, this data suggests a potential pathogenic role for THBS1, induced by the immune response and specifically driven by brain endothelial cells (Figure S6). This proof-of-concept analysis exemplifies how our generated data can add further knowledge on pathogenic mechanisms to disease associations found in clinical settings. Altogether, our findings illustrate synergistic pathogenic effects of *Pf-*PBMC with *P.falciparum-*iRBC, either through the induction of common pathways, such as JAK-STAT, or by driving transcriptional changes that further promote parasite sequestration.

## Discussion

Sequestration of *P. falciparum-*iRBC in the microvasculature has long been recognized as a key driver of CM pathogenesis, while the role of the immune response during pathogenesis remains poorly understood. Here, we used a human immunocompetent *in vitro* 3D-BBB model and demonstrate the effects of an immune-mediated mechanism of BBB disruption, acting independently of iRBC sequestration. We investigated this by stimulating PBMC from malaria-naïve donors with *P. falciparum* and perfusing them through a microfluidic, multicellular 3D-BBB model. Our analysis coupled single cell transcriptomics with functional barrier readouts and immunofluorescence validation. Our experimental approach aims to recapitulate immune signatures following the first encounter with the parasite. The development of protection against life-threatening malaria occurs quickly after the first exposure.^65–67^ Therefore, CM mostly affects children in regions of high malaria transmission intensity,^68^ or adults without prior malaria exposure,^69^ pointing towards the importance of studying the innate immune response to understand severe disease. This is further supported by controlled human malaria infection studies comparing European and African populations or reinfection studies, which reveal distinct immune signatures: naïve individuals show strong inflammatory and effector responses that are dampened and shift towards a regulatory response upon subsequent infections.^70–72^ Our analysis showed strong activation of CD14+ monocytes, NK cells and γδ T cells following *P. falciparum-*stimulation. The activation was marked by upregulation of inflammatory cytokines (TNF-α, IFN-γ) and cytotoxic effectors with vascular deleterious effects (granzyme B, perforin, Fas-L). Notably, this transcriptomic profile overlaps with gene modules that have been previously identified in severe malaria,^51,73–79^ highlighting that our *in vitro* stimulation recapitulates similar leukocyte activation signatures as observed during natural, severe *P. falciparum* infection.

The generation of an immunocompetent 3D-BBB model provides a significant step forward to understand the role of the immune response in CM, importantly providing functional readout to understand vascular damage mediated by human immune cells. Leukocyte accumulation in the brain microvasculature has been long identified in mouse ECM models^11,33^ and more recently in human postmortem samples.^8–10,80^ Our study demonstrates that *P. falciparum-*stimulation increases the binding capacity of specific PBMC populations, most prominently of activated CD8+ TEM and γδ T cell subpopulations. Notably, we show for the first time that *P. falciparum-*stimulation induces a shift towards higher levels of high-affinity LFA-1 conformation in T cells. The observed conformational transitions are quite likely induced by chemokine signaling, as similar mechanisms have been reported in response to chemokines^39,41^ or other pathogenic stimuli.^81^ Using ICAM-1 blocking antibodies, we validated LFA-1 - ICAM-1 as the dominant interaction mediating the adhesion of *P. falciparum-*stimulated leukocytes. This is consistent with a rodent ECM study which showed the capacity of LFA-1-blocking antibodies to prevent CD8+ T cell binding and disease development.^28,32^ Immune cell accumulation in malaria has been proposed to result from endothelial inflammatory activation, yet activation markers were not significantly elevated during the 1-hour perfusion time. Although endothelial activation is a well-established driver of leukocyte microvascular accumulation in malaria, our study illustrates that LFA-1 conformational activation constitutes an additional, early-acting mechanism that may be sustained throughout disease progression, and potentially exacerbated upon endothelial activation.

Postmortem samples have revealed strong endothelial inflammatory activation in the brain of CM patients, but it remains unknown whether this signature is driven mainly by the parasite or the immune response.^82^ Relatively short incubation periods with *Pf-*PBMC elicited strong inflammatory activation of all the BBB cells in the model. Although multiple cytokines were secreted by the stimulated leukocytes, we identified TNF-α and IFN-γ released by CD14+ monocytes, γδ T cells and NK cells as the main drivers for broad inflammatory activation in the 3D model, accounting for a remarkable fraction of differentially expressed genes associated with inflammation. This mirrors findings in humans, in which TNF-α and IFN-γ consistently emerge as important contributors to CM pathogenesis despite a diverse array of cytokines being elevated.^48,83,52,84,24^ Beyond driving inflammation, *P. falciparum-*stimulated leukocytes compromised BBB integrity by disrupting the endothelial cytoskeleton and inducing apoptosis, revealed independently by scRNA-seq analysis and confocal imaging. Barrier disruption strongly correlated with granzyme B and IFN-γ expression by γδ T cells and NK cells, whereas TNF-α secreted by monocytes appeared to act predominantly as an inflammatory driver rather than a mediator of barrier dysfunction. Consistent with our findings on the vascular disruptive effect of cytotoxic effectors, granzyme B-positive CD8+ T cells have been detected in brain postmortem samples^9^ and emerging clinical evidence points to a potential role for γδ T cells in severe malaria.^85^ Supporting this, mice lacking granzyme B and IFN-γ are protected from experimental CM.^50,86,87^ Strikingly, barrier disruption proved to be contact-dependent, as it was abrogated by inhibiting leukocyte-BBB adhesion with anti-ICAM-1 antibodies. This result fits with evidence that LFA-1 blockade prevents or reverses disease in mouse models of CM,^28,32^ highlighting a potential mechanism of leukocyte-binding dependent BBB disruption also in human disease. This requirement for direct contact aligns with the need for immunological synapse formation in granzyme B-mediated cytotoxicity,^88,89^ and with studies showing that IFN-γ activity declines markedly beyond ∼300 μm from its source.^90^ Our findings complement previous studies in 2D endothelial monolayers showing that NK cell secretion of IFN-γ after *P. falciparum* stimulation required direct endothelial contact,^91^ as well as an additional 2D study showing barrier disruption by *Pf-*PBMC.^92^ Altogether, our data indicate that *P. falciparum-*stimulated leukocytes drive BBB activation through cytokine secretion, while barrier disruption is linked to contact-dependent interactions and correlated with granzyme B and IFN-γ produced by γδ T cells and NK cells.

Collectively, these findings point towards potential benefits of immunomodulatory strategies in individuals lacking prior malaria exposure. Yet, such strategies will need to account for the concomitant vascular damage directly induced by the *P. falciparum* parasite. Our results highlighted overlapping pathogenic mechanisms between parasites and leukocytes, including perturbation of the JAK–STAT pathway. Ruxolitinib, a JAK inhibitor, has previously shown efficacy in preventing *P. falciparum*-induced barrier leakage in the 3D-BBB model.^18^ Furthermore, a controlled human malaria infection clinical trial has recently shown that Ruxolitinib prevents vascular damage and inflammation,^93^ highlighting JAK–STAT as a central, targetable pathway. While convergence on certain pathways is evident, parasites and immune responses also drive distinct signaling cascades that contribute independently to vascular pathology. More specifically, *P. falciparum-*stimulated leukocytes upregulated the expression of endothelial receptors, EPCR and ICAM-1, critical for parasite sequestration^55,94^ and strongly associated with CM.^56,57^ Conversely, a recent study demonstrated that *P. falciparum-*treated 3D brain microvessels triggered leukocyte recruitment.^14^ Together, these observations suggest a self-reinforcing loop that amplifies sequestration of both parasites and host immune cells at the vascular interface. Future studies should investigate the combined impact of *P. falciparum* and innate immune responses by sequential or simultaneous perfusion of these pathogenic drivers into the 3D-BBB model. Furthermore, we linked our results to a disease-associated haplotype at the population level, serving as a proof-of-concept how our data could be integrated with clinical association studies. The use of a highly tunable, bioengineered model, in combination with as a series of experimental and computational tools, has allowed us to unravel the complex immune-vascular interactions and molecular mechanisms leading to BBB damage in CM, suggesting novel therapeutic strategies.

### Limitations of the study

The difficulties to access the brain in CM patients have hampered our understanding of the disease. While our study overcomes the inter-species differences of rodent ECM, *in vitro* models entail inherent limitations. Our analysis used PBMC stimulated with uninfected RBC as a negative control to reveal specific responses elicited by *P. falciparum-*stimulation, yet the possibility that an endothelial-immune mismatch between donors might promote endothelial disruption was considered. The lack of BBB inflammatory activation and preserved permeability baseline values in samples exposed to control PBMC excluded a direct mismatch effect and the short co-culture time rendered unlikely the development of a specific allogeneic T cell response. Moreover, the response of NK cells from a selected donor bearing a KIR genotype matched to endothelial HLA class I allotypes was comparable to the effect observed with different samples, excluding that NK cell alloreactivity boosted following *P. falciparum-*stimulation explained the vascular damage. As an alternative, iPSC-based models or access to donor-matched immune and BBB samples would in principle allow the possibility of future isogenic models. A second limitation is that we did not include responses from other immune cells, such as neutrophils or platelets in our experiments. Nevertheless, future iterations of the model could include these cell types. One final limitation is the focus on the leukocyte response of naïve donors. While this models CM disease occurring in malaria-naïve adults, CM often affects pediatric populations in regions with high malaria endemicity. Future studies would need to assess the immune response in pediatric samples from endemic populations to address potential geographical and age-mediated differences in immune responses, as well as to identify immune signatures caused by prior exposure to the parasite.

## Material and Methods

### Primary human cell culture

Primary human brain microvascular endothelial cells (HBMEC, Cell Systems) were cultured in microvascular endothelial cell growth medium-2 (EGM-2MV, Lonza) up to passage 8. Primary human astrocytes (HA, ScienCell) were maintained in basal media supplemented with 2% FBS, 1% Pen-Strep solution, and 1% astrocyte growth supplement (ScienCell, #1852) up to passage 7. Primary human brain vascular pericytes (HBVP, ScienCell) were cultured in basal media containing 2% FBS, 1% Pen-Strep solution, and 1% pericyte growth supplement (ScienCell, #1252) up to passage 8.

### 3D-BBB model fabrication

3D-BBB devices were fabricated as previously described.^18,26^ Briefly, type I collagen was extracted from rat tails and dissolved in 0.1% acetic acid to a stock concentration of 15 mg/mL. The stock collagen was neutralized with sodium hydroxide (NaOH) and M199 (Gibco) and diluted to 7.5 mg/mL with EGM-2MV supplemented with 1% astrocyte and pericyte growth factors (ScienCell). Primary HA and HBVP were added to the neutralized collagen solution in a 7:3 ratio, using a concentration of 7.5×10^5^ HA /mL : 3.2×10^5^ HBVP /mL in the collagen solution. Soft lithography and injection molding was used to fabricate the 3D-BBB model.^26^ A network of micro-patterned channels was fabricated through soft-lithography by injecting a collagen hydrogel through an injection port between a top polymeric jig and a positive polydimethylsiloxane (PDMS) pre-patterned mold, previously functionalized with a O^2^ plasma treater. We used a 13 by 13 channel grid pattern creating 120 μm diameter microvessels that recapitulates a wide gradient of flow mechanical cues in a single device. The bottom part was fabricated by spreading the collagen solution on top of a coverslip within the bottom housing jig and by flattening it using a flat PDMS stamp to obtain a thin collagen layer. After 75-minute 37°C collagen gelation, the PDMS stamps were removed, and the top and bottom sections were assembled. The 3D-BBB devices were incubated with EGM-2MV medium supplemented with 1% astrocyte and pericyte growth factors (ScienCell) overnight before HBMEC seeding. Primary HBMEC were seeded at a concentration of 7×10^6^ cells /mL through gravity-driven flow, adding 8 µL cell suspension to the device in multiple seeding rounds until full coverage of the microfluidic network was reached. 3D-BBB models were cultured for up to 6 days and media was changed every 12 hours by gravity-driven flow.

### P. falciparum culture

HB3var03 variant *P. falciparum* parasites were cultured in human 0+ erythrocytes in RPMI 1640 medium (Gibco) supplemented with 10% human type AB-positive plasma, 5 mM glucose (Sigma), 0.4 mM hypoxantine (Sigma), 26.8 mM sodium bicarbonate (Corning/Sigma) and 1.5 g/L gentamicin (Gibco). Plasma and 0+ red blood cells were obtained through a local collaboration agreement with the Catalan Blood Bank. *P. falciparum-*iRBC were grown in sealed top T75 flasks in a gas mixture of 90% N^2^, 5% CO^2^ and 1% O^2^. Culture was maintained at a parasitemia between 0.5% and 8% and at 4% hematocrit.

For PBMC stimulation, parasites were regularly synchronized in their ring stage with 5% sorbitol (Sigma) or in their schizont stage using a 70% Percoll gradient (Cytiva). Subsequently, parasites at 36-44 hours post-infection were purified by magnetic activated cell sorting (MACS) using MACS LD columns (Miltenyi Biotech), achieving a purity of 85-99% mature schizonts.

### PBMC culture and stimulation with *P. falciparum* iRBC- or RBC

PBMC from healthy European/US adult donors living in non-malaria endemic regions with blood group 0+ were purchased from StemCell Technologies. Donor consent and ethical compliance were managed by the vendor. Donors were balanced with respect to ethnicity (Caucasian vs. African American) and sex (male vs. female). PBMC from one additional donor were locally extracted and were HLA-matched with respect to the NK cell Killer Cell Immunoglobulin-like Receptor (KIR) to the endothelial cells. Written informed consent was obtained, and the study protocol was approved by the institutional ethics committee (Clinical Research Ethics Committee, Parc de Salut Mar CEIC 2018/7873/I).

Two days before 3D-BBB device perfusion, PBMC were thawed in RPMI media (RPMI 1640, Sigma), supplemented with 10% heat-inactivated fetal bovine serum (Gibco), 0.5% penicillin/streptomycin (Gibco), and 1% Glutamax (Gibco). PBMC were washed once in media and allowed to rest for 24 hours at 37°C with 5% CO^2^ at a cell density of 2-4 x 10⁶ cells per mL in a 50 mL Falcon tube with a loose lid for proper gas exchange. On the following day, PBMC were plated at 5×10^5^ cells/well in a 96-well round-bottom plate in RPMI / 10% FBS media and were stimulated with mature schizonts (36-44 hours post-infection) at a 1:3 ratio (1.5×10^6^ iRBC /well), ensuring complete egress of all iRBC during the incubation time. For the control condition, PBMC at the same density were incubated at a 1:3 ratio with RBC (1.5×10^6^ RBC /well), prepared as described above. PBMC were incubated with iRBC or RBC for 16 hours at 37°C, ensuring schizont rupture after 4-12 hours. When required for intracellular cytokine flow cytometry analysis, brefeldin A (Sigma) was added for the last 12 hours at a final concentration of 10 μg/mL.

### Sample incubation with *P. falciparum*-stimulated or control PBMC

3D-BBB devices were perfused after 6 days in culture, the timepoint in which they present peak barrier function.^18^ Stimulated PBMC were resuspended in coculture media (RPMI 1640 (Sigma) supplemented with 5% heat-inactivated fetal bovine serum (Gibco), 0.4% recombinant human FGF (Lonza), 0.1% vascular endothelial growth factor (Lonza), 0.1% R3-insulin-like growth factor (Lonza), 0.1% ascorbic acid (Lonza), 0.1% recombinant human endothelial growth factor (Lonza), 0.5% penicillin/streptomycin (Gibco), 1% Glutamax (Gibco) and 1% astrocyte and pericyte growth supplement (ScienCell) to remove any soluble iRBC remnants at a concentration of 3 x 10^6^ PBMC per ml. 3D-BBB models were perfused through gravity-driven flow for 1 hour by removing and adding fresh cell suspension every 10 minutes, which represented a total perfusion of roughly 1-1.2 x 10^6^ PBMC. After perfusion, devices were washed for 15 minutes with media and incubated for 10 hours at 37°C.

For high content imaging analysis, HBMEC were seeded on poly-L-lysine coated 96-well plates (PhenoPlate 96-well, PerkinElmer, 6055300) at density of 10^4^ cells per well and grown until reaching confluency. Monolayers were incubated with 1×10^5^ control or *Pf-*PBMC per well in 150 µl coculture media (described above). As a control, monolayers were incubated with the equivalent concentration of iRBC remnants (3×10^5^ iRBC per well) spun down and resuspended in the same way as iRBC-stimulated PBMC.

### Immunofluorescent staining of 3D-BBB models

After 10 hours of incubation with perfused, cytoadherent PBMC, 3D-BBB devices were fixed with ice-cold 4% paraformaldehyde (PFA) by gravity-driven flow for 20 minutes. The devices were washed three times for 10 minutes with phosphate-buffered saline (PBS) before immunostaining. To prevent collagen-related background signal 3D-BBB devices were treated with background buster (Innovex Biosciences) for 30 minutes. For CD3 and CD45 staining, blocking and permeabilization was performed with 2% bovine serum albumin (BSA) and 0.1% Triton-X-100 (in PBS) for 5 min followed by perfusion of 2% BSA and 0.1% Tween (in PBS) for one hour at room temperature. For all other antibodies only 2% BSA and 0.1% Triton X-100 (in PBS) was applied for one hour. The following primary antibodies were added in 2% BSA in PBS and incubated overnight: CD3-Alexa Fluor 594 (R&D Systems, FAB100T, 1:200), CD3-Alexa Fluor 555 (Abcam, ab208514, 1:200), CD45-Alexa Fluor phycoerythrin (Biolegend, 304008, 1:200), CD45-Alexa Fluor 488 (Abcam, ab200315, 1:200). Primary antibodies against ICAM-1 (Abcam, ab20, 1:100), PDGFRβ (Abcam, ab69506, 1:100), VE-cadherin (Santa Cruz, sc-52751, 1:100), VE-cadherin (Abcam, ab33168, 1:100), GFAP (Abcam, ab4674, 1:200), cleaved caspase 3 (Cell Signaling, 9661S, 1:100), and phalloidin-594 (ThermoFisher, A12381, 1:100) were incubated overnight in 2% BSA and 0.1% Triton X-100. After three 10-minute PBS washes, the devices were incubated with DAPI (ThermoFisher D21490, 8 μg/mL), and Alexa-Fluor 488-, Alexa-Fluor 594-, Alexa-Fluor 561-, or Alexa-Fluor 647-conjugated secondary antibodies (ThermoFisher, 1:250) for 1 h at room temperature followed by three more washes with PBS.

### Immunofluorescent staining of 2D monolayers

HBMEC 2D monolayers were fixed with ice-cold 4% PFA for 20 minutes and washed three times with PBS. Blocking and permeabilization were performed for one hour using 2% BSA and 0.1% Triton X-100 (Sigma) in PBS at room temperature. Primary antibodies against ICAM-1 (Abcam, ab20, 1:200) were diluted in 2% BSA and 0.1% Triton X-100 in PBS and incubated for one hour. Monolayers were washed three times with PBS, incubated in 2% BSA and 5% neutralizing goat serum (Gibco) with DAPI (ThermoFisher, D21490, 8 μg/mL), Alexa-Fluor 488-, Alexa-Fluor 594-, or Alexa-Fluor 647-conjugated secondary antibodies (ThermoFisher, 1:250) for one hour at room temperature before washing three times with PBS.

### Image acquisition and analysis

2D monolayers were imaged using an Opera Phenix HSC system (PerkinElmer) and processed with a custom python script (combining z-stacks and channels) and Fiji (ImageJ, v1.54p). 3D-BBB devices were imaged using a LSM980 Airyscan 2 microscope (Zeiss) or a Thunder Imager Live Cell & 3D Assay (Leica) (whole device overviews) and processed using Fiji (ImageJ, v1.54p). For the quantification of PBMC binding to the 3D-BBB model, images of the bottom section of the outermost grid channel were acquired, representing a gradient of wall shear stress between the corners and inlet/outlet. Maximum intensity projections of stacks with 5 μm distance were performed. Thresholds using the *Otsu* method were applied specific to each edge and staining, followed by dilation and filling holes operations. Fiji Image Calculator was used to perform arithmetic operations between the channel masks, generating binary masks of overlapping fluorescence signals for nuclei (DAPI) and PBMC markers (CD45 and CD3). Eleven rectangular regions of interest (ROIs) with dimensions of 350 x 80 μm were selected along the channel keeping a 10μm distance from the channel edges to restrict analysis to the central part of the channel where flow remains laminar (Figure S2E). The eleven chosen ROI represent the outer edges of the 13 x 13 grid, excluding the inlet and outlet areas. Double- or triple positive signals of 10-60 μm^2^ in size were automatically counted in the selected ROI.

For the analysis of ICAM-1, Phalloidin, cleaved caspase 3, and VE-cadherin staining, images were taken in ROIs corresponding to 5 channel junctions distributed throughout the 3D-BBB model in areas of different wall shear stress, and maximum intensity projections of z stacks were generated. To measure ICAM-1 intensity, the mean grey value per pixel was determined. For actin stress fiber quantification phalloidin fluorescence intensity over the endothelial nucleus was quantified as follows: A mask of endothelial nuclei was created and the mean phalloidin intensity within the nuclear mask was normalized by the total phalloidin intensity throughout the entire image. Cleaved caspase 3 signal was counted manually in endothelial cells and the ratio of positive cells was calculated using automated segmentation of endothelial nuclei. Junctional VE-cadherin was quantified by applying a Tophat filter (rolling ball radius = 10 μm), followed by Otsu thresholding and measuring the area of the generated mask per endothelial cell.

### Barrier integrity xCELLigence assay

xCELLigence 96 well PET E-plates (Agilent, 300600910) were coated with poly-L-lysine and HBMEC were plated at a density of 10^4^ cells. Baseline cell growth measurements were performed for up to 48 hours until cell index values plateaued indicating a confluent monolayer. 10^5^ control or *Pf-*PBMC per well were added in 150 µl coculture media (described above). For transwell experiments, xCELLigence E-plate inserts (Agilent, 6465412001, 0.4µm) were used. After baseline measurements, media was added to the wells, followed by placing the transwell inserts and filling them with PBMC cell suspension. Subsequently, the cell index was measured in 15-second intervals for 12 hours and then every 15 minutes for another 26 hours. All conditions were measured in duplicates and normalized to a media control as follows: The timepoint before PBMC addition was subtracted from all following timepoints and all conditions were normalized to the media control cell index by subtraction. The area under the curve (AUC) was quantified by taking the integral of the negative portion of the cell index plot, indicative of barrier disruption for a timeframe of 38 hours after PBMC addition.

### Microvascular permeability assays

Microvascular permeability was determined as previously described.^18^ Permeability assays were performed on the 3D-BBB model at day 6 after fabrication. Using a withdrawing syringe pump (Harvard Apparatus PHD ULTRA) 70 kDa FITC-dextran at a concentration of 100 µg/mL was perfused at a flow rate of 10 µL/min. Over a timeframe of 10 minutes confocal images were acquired every 2.5 minutes with 5 μm z-step size in 5 different areas within the 3D-BBB model. Permeability measurements on the same areas were performed at timepoint 0h and at timepoint 10h after perfusion with *Pf*-PBMC or control PBMC. Equally sized regions of interest (ROIs) (250 µm x 150 µm) were selected along the microvessel wall to include a specified area of the vessel (250 µm x 30 µm) and of the collagen hydrogel (250 µm x 120 µm). For each ROI, fluorescence intensities were measured at time point t0 (after complete vessel filling with dextran) and t1 (2.5 minutes after). The microvascular apparent permeability was quantified using the following formula:

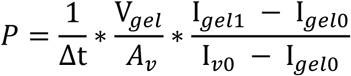

Δt is the time interval between the two frames t0 and t1, V_gel_ is the volume of the collagen matrix, A_v_ is the lateral vessel surface, I_gel1_ - I_gel0_ is the difference in fluorescence intensities in the gel area between t1 and t0, and I_v0_ is the fluorescence intensity in the vessel at t0. Permeability was calculated before and after PBMC incubation, respectively, and a permeability ratio was calculated for each ROI.

### Flow cytometry assay

After 16 hours of PBMC stimulation, antibodies specific for high-affinity conformations of LFA-1 (LFA1-PE/Cy7 m24, Biolegend, 363417) and VLA-4 (VLA4-PE HUTS-21, BD Biosciences, 556049) were added to the media for 15 minutes followed by EDTA (Invitrogen) for 10 minutes. Subsequently, blocking was performed using a solution of human IgG (Baxter, 812651, 10 µg/mL) including the LiveDead^TM^ Fixable Near-IR dead cell stain kit (ThermoFisher, L34975) to assess viability. PBMC were surface stained using the following monoclonal antibodies: CD3-PerCP (Biolegend, 981016), CD56-BV605 (BD Biosciences, 562780), CD19-FITC (BD Biosciences, 555412), CD14-APC (eBiosciences, 17-0149-42), and TCRγ/δ1-PE/Cy7 (BD Biosciences, 655410). For intracellular staining, cells were fixed and permeabilized (BD Cytofix/Cytoperm, BD Biosciences, 554722), according to manufacturer instructions, followed by intracellular staining with TNF-α-PacificBlue (Biolegend, 502920) and IFN-γ-PE (BD Biosciences, 554701). Data were acquired on an LSR Fortessa flow cytometer (BD Biosciences) and analyzed using FlowJo software (v10.9.0).

### Luminex assay

Secreted protein concentrations were measured in the supernatants of PBMC from 6 different donors after 16 hours of iRBC- or uninfected RBC-stimulation. We quantified 25 cytokines, chemokines, growth factors, and related biomarkers using a customized Human Magnetic 25-plex Panel (Bio-Techne R&D Systems) on a Luminex® 100/200 platform at the Immune Response and Biomarkers Core Facility at ISGlobal. The panel included: Granzyme B, IFN-α, IFN-γ, IL-1β, IL-2, IL-6, IL-10, IL-12p70, IL-15, IL-18, matrix metalloproteinase-1 (MMP-1), TNF-α, vascular endothelial growth factor A (VEGF-A), Angiopoietin-2, monocyte chemoattractant protein-1 (MCP-1/CCL2), macrophage inflammatory protein-1α (MIP-1α/CCL3), MIP-3β, RANTES (CCL5), eotaxin, 6Ckine/CCL21, monokine induced by IFN–γ (MIG/CXCL9), IFN-γ induced protein-10 (IP-10/CXCL10), Interferon-inducible T cell alpha chemoattractant (I-TAC/CXCL11), Stromal cell-derived factor-1α (SDF-1α/CXCL12), and triggering receptor expressed on myeloid cells 1 (TREM-1). The assay included 16 serial 2-fold dilutions of a standard with known analyte concentrations and two blank controls. Following the manufacturer’s protocol, 50 μL of each sample or standard was incubated with the multiplexed magnetic beads for 2 hours at room temperature on a shaker at 800 rpm. After washing, beads were incubated with a biotinylated antibody cocktail for 1 hour at room temperature, followed by a 30-minute incubation with streptavidin-PE under the same conditions. Beads were then washed, resuspended in 100 μL of wash buffer, and acquired using xPONENT® software version 3.1. Analyte concentrations were determined by interpolating blank-subtracted median fluorescence intensity (MFI) values against a 5-parameter logistic standard curve and expressed in pg/mL. Values below the lower limit of detection (mean of blanks + 2 standard deviations) were imputed as half of the lowest detected analyte concentration, while those exceeding the upper limit of quantification were assigned double of the highest detected analyte concentration. CXCL12, CCL11, IL-12p70 were excluded from the downstream analysis due to too many values below the detection limit.

### Sample preparation for single-cell RNA sequencing

After the 16-hour stimulation, unperfused PBMC were resuspended in RBC lysis buffer (eBiosciences) and incubated for 5 minutes at room temperature. The reaction was stopped by adding 6 ml phosphate buffered saline (PBS) and cells were washed two times at 4° C in PBS containing 0.1% bovine serum albumin (BSA). The single-cell suspension was filtered through a 35 µm cell strainer (Falcon) and cells were counted and concentration adjusted for 10x Genomics barcoding.

To generate the perfused dataset, the 3D-BBB models were fabricated, perfused, and incubated as described above. Three devices per condition were disassembled, and the middle, microfluidic pattern-containing collagen part was dissected. The collagen piece was dissociated for 8 - 10 minutes (until complete dissociation) in collagenase diluted in serum-free media (1 mg/mL, Sigma). The collagenase reaction was stopped with complete media and the cells were trypsinized for 8 minutes to obtain a single-cell solution. The cells were mechanically dissociated by repeated pipetting with wide-bore tips and washed two times in PBS containing 0.1% BSA. The single-cell suspension was filtered through a 35 µm cell strainer (Falcon, 352235) and cells were counted and concentration adjusted for 10x Genomics barcoding.

### Single-cell RNA barcoding and sequencing

mRNA transcripts of each cell were barcoded using the Chromium Controller (10x Genomics, firmware version 4.00). The reagent system was Chromium Single Cell 3′ GEM, Library & Gel Bead Kit v3.1 (10x Genomics) and a Chromium Next GEM Chip G Single Cell Kit (10x Genomics). Barcoding and cDNA library construction were performed according to the manufacturer’s instructions. Finished cDNA libraries were sequenced with NextSeq2000 (Illumina). We read 28 base pairs (bp) for 10x Genomics barcodes with unique molecular identifiers (UMIs), 10 bp for i7 and i5 indices, and 90 bp for fragmented cDNA.

### Single-cell RNA sequencing data analysis

#### Sequence alignment

Sequenced reads were aligned to a combined reference genome constructed from the human genome (GRCh38) and the *P. falciparum* genome (hb3) to generate the feature barcode matrices with the *CellRanger* pipeline (v. 7.0.1, 10x Genomics). All downstream data analysis was performed with R version 4.4.1^95^ or Python version 3.12.9.^96^

#### Quality Control (QC)

Quality control was performed using the *scuttle* package (v.1.14.0)^97^ as follows: Genes that were found in less than 2 cells in an experimental dataset were excluded. The QC thresholds were identified based on scatter plots of detected gene counts against the proportion of mitochondrial gene expression in each cell and only keeping the cells above and below the determined detected gene cutoffs (PBMC datasets: lower cutoff: 800-1400 / higher cutoff: 5000, PBMC-perfused 3D-BBB model datasets: lower cutoff: 350-700) and below the mitochondrial gene cutoff (PBMC datasets: 6% - 8%, PBMC-perfused 3D-BBB model datasets: 6.5% - 7.5%) for further analysis. Subsequently, doublets were excluded using the *scDblFinder* package (v. 1.18.0).^98^

#### Data normalization

The raw UMI counts of the QC-filtered cells were normalized using the deconvolution approach from the R package *scran* (v.1.32.0).^99^ The size factor for the library size correction of each cell was calculated with the *calculateSumFactors* function and the raw counts were normalized based on the size factor and log2-transformed with the *logNormCounts* function. These values appear as “logcounts”.

#### Dataset integration

To account for different sequencing depths per-batch scaling normalization was performed on the three sequencing datasets from different PBMC donors using the *multiBatchNorm* function in the package *batchelor* (v.1.20.0).^100^ The datasets were integrated using the *Seurat* (v.5.1.0)^101^ canonical correlation analysis (CCA) approach. Therefore, each dataset was normalized using the *NormalizeData* function and highly variable genes were selected using the *FindVariableFeatures* function, followed by scaling with the *ScaleData* function and principal component analysis with the *RunPCA* function. Subsequently, CCA integration was performed using the *IntegrateLayers* function and the corrected results were used for Uniform Manifold Approximation and Projection (UMAP) visualizations, while the non-corrected counts and logcounts matrices was used for differential gene expression and all other analyses.

#### UMAP visualization and clustering

After normalization and dataset integration, the UMAP was constructed from the newly calculated, low-dimensional aligned canonical correlation vectors (*integrated.cca*). Cell population clusters were identified using the *bluster* package (v.1.14.0)^102^ and the *Leiden* algorithm.^103^ Cell type assignment was performed manually based on the cluster marker genes. Due to low cell numbers, annotation of adherent immune cell subpopulations in the PBMC-perfused 3D-BBB datasets was performed via label transfer from the unperfused-PBMC dataset using the Seurat *FindTransferAnchors* and *TransferData* functions.

#### Differential expression analysis

Differential expression analysis was performed separately for each cell type utilizing the hurdle (two-part generalized regression) model from the *MAST* package (v.1.30.0).^104^ The Benjamini-Hochberg method was applied to the p-values to account for multiple testing. All significantly differentially expressed genes for all cell types and conditions can be found in Supplemental Table S1. The heatmaps visualizing the log2-transformed fold change (log2FC) values were created using *pheatmap* (v.1.0.12).^105^

#### Gene ontology (GO) term over-representation analysis

GO-term over-representation analysis was performed on significantly upregulated genes shared between endothelial cells and pericytes (FDR < 0.05, log2FC > 0.05) using the *enrichGO* function (Benjamini-Hochberg correction, p-value cutoff 0.05, max. geneset size 2000) from the *clusterprofiler* package (v.4.12.0),^106^ including “Biological Process” GO-terms. The *pairwise_termsim* and *emapplot* functions from the *GOSemSim* package (v.2.30.0)^107^ and the *enrichplot* package (v.1.24.0)^108^ were used to visualize the analysis results. GO-term clusters in the enrichment map were manually labeled. The list of all significant GO-terms can be found in Supplemental Table S2.

#### Human Molecular Signatures Database (MSigDB) hallmark gene set over-representation analysis

MSigDB hallmark gene set over-representation analysis was performed on significantly upregulated genes in each immune cell type (FDR < 0.05, log2FC > 0.2) using the *msigdbr* package (v.7.5.1)^109^ and the *compareCluster* function (fun = “enricher”, Benjamini-Hochberg correction, p-value cutoff 0.05, max. geneset size 1500) from the *clusterprofiler* package. Hallmark gene sets significant in at least 2 cell types are shown and all significant gene sets can be found in Supplemental Table S2.

#### Comparative GO term over-representation analysis including iRBC-egress dataset

For comparative GO-term over-representation analysis the presented scRNA-seq data was compared with a previously published dataset.^18^ GO-term over-representation analysis was performed on significantly upregulated genes in endothelial cells upon exposure to iRBC-egress or *Pf*-PBMC (FDR < 0.05, log2FC > 0.05) using the *compareCluster* function (fun = “enrichGO, Benjamini-Hochberg correction, p-value cutoff 0.05, max. geneset size 500) from the *clusterprofiler* package including all GO-term categories. The *pairwise_termsim* and *emapplot* functions from the *GOSemSim* package and the *enrichplot* package were used to visualize the analysis results. GO-term clusters in the enrichment map were manually labeled. The list of all significant GO-terms can be found in Supplemental Table S2.

#### Gene signature score calculations

Gene signature scores were calculated using the *AddModuleScore* function from *Seurat*. Gene signature scores were created from gene lists as follows:

*T cell cytotoxic and effector signature* (Figure 3B): *TNF, IFNG, PRF1, GZMB, CD69, TNFRSF9, LAG3, TNFRSF4, CRTAM, NKG7*

*Endothelial & pericyte shared inflammatory gene set* (Figures 2C and S4F): significantly upregulated genes shared between HBMEC and pericytes (FDR < 0.05, log2FC > 0.05)

*Disruptive module signature* (Figures 6E and S5F): top 100 hub genes sorted by eigengene-based connectivity (kME) of module 1, 5, and 6, respectively

#### Gene co-expression network analysis

The *hdWGCNA* package (v. 0.4.5)^43^ was used to perform weighted gene co-expression network analysis (WGCNA) following the basic vignette workflow as follows: The dataset was set up for WGCNA by selecting genes that are expressed in at least 5% of cells. Metacells were constructed by grouping cells by dataset and cell type and using k = 25 as k-nearest-neighbors parameter and subsequently normalized. The co-expression network was constructed using endothelial cells as the cell type of interest using the automatically selected soft power. Module eigengenes were computed as per standard settings and harmonized by dataset (hME) followed by calculation of eigengene-based connectivity (kME) values representing module “hub genes”. hME were used for UMAP and violin plot visualizations. GO-term over-representation analysis was performed top 100 hub genes sorted by kME of module 1, 5, and 6, respectively, using the *enrichGO* function (Benjamini-Hochberg correction, p-value cutoff 0.05, max. geneset size 2000) from the clusterprofiler package including “Biological Process” GO-terms. A Mann Whitney U test with Bonferroni multiple testing correction was performed to identify differentially expressed modules between the conditions (adjusted p-value < 0.05, expression in > 30% of cells, log2FC > 2).

#### Ligand-receptor interaction analysis

Ligand-receptor interaction analysis was performed using the *CellChat* package (v.2.1.2).^21^ Cell types with less than 150 cells were excluded from the analysis and the datasets were filtered to only the *Pf*-PBMC condition. CellChat was run on either the unperfused *Pf-*PBMC or the *Pf-*PBMC-perfused 3D-BBB model datasets using 20% truncated mean to calculate the average gene expression per cell type. The inferred communications were subset to the pathways of interest for unperfused *Pf-*PBMC analysis (*CXCL, CCL*) (Figure 3C and Figure S3C) or *Pf-*PBMC-perfused 3D-BBB model analysis (*ICAM, VCAM, MHC-I, SELE, SELL, MHC-II, CXCL, CCL*) (Figure 6D), respectively.

For differential ligand-receptor analysis, *CellChat* was run on cells from either of the two experimental conditions separately before the *CellChat* objects were merged and the interactions compared. Up and down-regulated ligand-receptor signaling pairs were identified from the differential expression analysis using the *identifyOverExpressedGenes* function (thresh.fc = 0.1, thresh.p = 0.05). The Chord diagram was constructed and strongest ligand-receptor interactions are labeled (Figure S1D).

#### Differential binding analysis (scCODA)

Differential binding analysis for immune cell types and T cell subpopulations was performed separately, using the Bayesian model from the *scCODA* package (v. 0.1.9).^110^ The compositional model was set up using “condition + dataset” as the formula to account for the paired design and the reference cell type was selected automatically.

#### Cytokine-target analysis

The ligand–target regulatory potential matrix was derived from the *nichenetr* package (v.2.2.0)^53^ (https://zenodo.org/record/7074291/files/ligand_target_matrix_nsga2r_final.rds). For the cytokine-target flow diagram, this matrix was filtered for cytokines previously identified with significantly upregulated levels in *Pf*-PBMC from the Luminex protein quantification and significantly upregulated (target) genes in endothelial cells (FDR < 0.05, log2FC > 0.1). The matrix was binarized using a regulatory potential cutoff of 0.2 only keeping cytokines and target genes with regulatory interactions above this threshold. Immune cell types were considered contributors to the increased cytokine levels if the cytokine was significantly upregulated in the respective cell type (FDR < 0.05, log2FC > 0.05). For cytokines that were not significantly upregulated in any individual cell type, cell types were ranked by their total cytokine expression. The top cell types cumulatively contributing to 50% of the total cytokine expression were then considered as contributors for that specific cytokine. The flow diagram was generated using the *ggsankey* package.^111^ To evaluate the contribution of cytokines to endothelial differential gene expression, the percentage of significantly upregulated endothelial genes with a cytokine among their top-5 regulatory ligands was determined among all significantly upregulated genes (FDR < 0.05, log2FC > 0.05) in endothelial cells.

### Statistical analysis

Statistical analysis was performed using R (v. 4.4.1). Two-tailed Mann Whitney U test was used to analyze non-normally distributed samples. Multiple conditions were analyzed using a Kruskal-Wallis test followed by Dunn’s test with Benjamini-Hochberg multiple testing correction. Paired samples were analyzed using a paired two-tailed Student’s test with Benjamini-Hochberg multiple testing correction. Statistical significance was defined as p < 0.05 unless otherwise specified. Values are presented as the median with the interquartile range (IQR).

## Lead contact

Further information and requests for resources and reagents should be directed to and will be fulfilled by the lead contact, Maria Bernabeu.

## Data and code availability

Data availability: due to patient consent and confidentiality agreements, raw sequencing datasets can be accessed for validation or reuse by contacting the lead contact. The raw scRNA-seq count tables from all experiments can be found in <u>E-MTAB-15663</u> and <u>E-MTAB-15664</u>.

Code availability: all code necessary for the data analysis and visualization will be available at https://github.com/Alina-Ba/scRNAseq_Pf-PBMC. The private link can be shared with the reviewers upon request. Any additional information required to reanalyze the data reported in this paper is available from the lead contact upon request.

## Acknowledgements

We would like to thank all members of the Bernabeu lab as well as Chris Moxon for supportive discussions and critical feedback. We want to thank Sergi Beneyto Calabuig and Lars Velten for their input on the scRNA-seq data analysis. We thank Nicola Gritti and the EMBL Mesoscopic Imaging Facility (MIF) for discussions on image analysis support and scripts. We are grateful to the Genomics Core (GeneCore) facility at EMBL, Genomics Core facility at the Universitat Pompeu Fabra (UPF), the Genome Biology Computational Support (GBCS) (Charles Giradot) at European Molecular Biology Laboratory (EMBL) for their sequencing support and to EMBL IT Service for the service of high-performance computing resources. We thank Ruth Aguilar from Immune Response and Biomarkers Core Facility at ISGlobal running the Luminex assay and her data analysis input and the EMBL Bio-IT Centre for Statistical Data Analysis for their data analysis support. We thank Marina Fortea Guillamon and Joana Rafaela Mendonca da Silva for imaging assistance. We are grateful to the UPF Flow Cytometry Core Facility and the EMBL Flow Cytometry Core Facility for their support and analysis infrastructure. We thank Dr. Carlos Vilches (previously working at Immunogenetics and Histocompatibility Lab, Instituto de Investigación Sanitaria Puerta de Hierro - Segovia de Arana, Madrid, Spain) for performing HLA-typing of our primary cells. The great majority of this work was funded by the European Research Council (ERC) under the European Union’s Horizon 2020 research and innovation program (Grant agreement no. 948088), the EMBL core program funding and contributions from the EMBL Infection Biology Transversal Theme. H.F is supported by a fellowship from the EMBL Interdisciplinary (EIPOD4) program under Marie Skłodowska-Curie Actions COFUND (847543), and O.R.O by an EMBL EIPOD-LinC fellowship. GM was supported by RYC 2020-029886-I/AEI/10.13039/501100011033, co-funded by European Social Fund (ESF).

## Use of AI tools

During the preparation of this work the author(s) used ChatGPT 5 in order to assist with main manuscript grammar and phrasing, as well as coding suggestions. After using this tool/service, the author(s) reviewed and edited the content as needed and take(s) full responsibility for the content of the published article.

## Author contributions

A.B and M.B conceived the work.

A.B, P.D, F.N, M.A, H.F, L.P., G.M and M.B designed experiments.

A.B, P.D, F.N, M.A, H.F, L.P. performed experiments with assistance of S.S, M.F, O.R.O, B.L.G A.B, P.D, M.A analyzed the experimental data under the guidance of M.L.B and M.B.

A.B and F.N performed the scRNA-seq experiments with input from D.S on experimental planning, under the guidance of L.M.S, J.S and M.B.

A.B, J.B, A.R.G, F.N. performed the scRNA-seq data analysis under the guidance of L.M.S, J.S and M.B.

G.M. provided critical conceptual input.

A.B and M.B wrote the original draft of the manuscript. All authors contributed to manuscript writing, revision, editing and suggestions.

M.B contributed to project supervision, administration and funding acquisition.

