## Supplemental information for "Innate immune responses to *Plasmodium falciparum* disrupt the blood-brain barrier"

[1] European Molecular Biology Laboratory (EMBL) Barcelona, Barcelona, Spain
[2] Collaboration for joint PhD degree between EMBL and Heidelberg University, Faculty of Biosciences, Heidelberg, Germany
[3] Humboldt University, Faculty of Life Sciences, Berlin, Germany
[4] Department of Genetics, Stanford University School of Medicine, Stanford, CA, USA.
[5] University Pompeu Fabra, Barcelona, Spain
[6] European Molecular Biology Laboratory (EMBL), Cell Biology and Biophysics Unit, Heidelberg, Germany
[7] Hospital del Mar Research Institute, Barcelona, Spain
[8] Institució Catalana de Recerca i Estudis Avançats (ICREA), Barcelona, Spain
[9] European Molecular Biology Laboratory (EMBL), Genome Biology Unit, Heidelberg, Germany
[10] Stanford Genome Technology Center, Palo Alto, CA, USA
[11] ISGlobal, Barcelona, Catalonia, Spain
[12] CIBER Enfermedades Infecciosas (CIBERINFEC), Instituto de Salud Carlos III, Barcelona, Spain

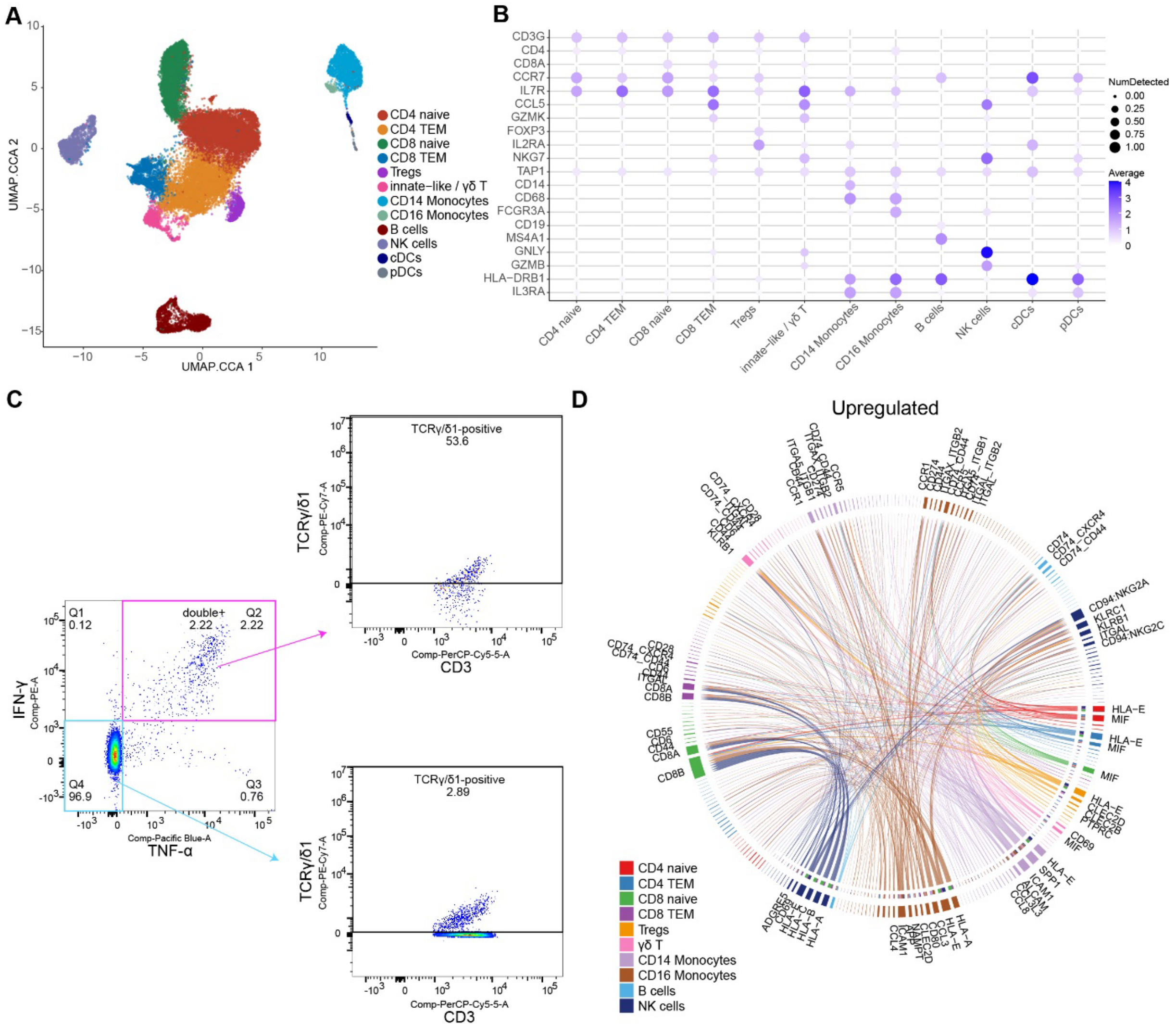


**Figure S1. Characterization of *P. falciparum*-stimulated PBMC.**

**A.** UMAP visualization of sequenced PBMC, colored by cell type annotation, including identified subpopulations.

**B.** Dot plot of canonical PBMC cell type markers.

**C.** Representative intracellular cytokine analysis by FACS of *P. falciparum*-stimulated T cells. Among activated, double-positive (TNF-α-positive and IFN-γ-positive) T cells, 53.8% express the γδ T cell marker TCRγ/δ1, compared to 2.89% of double-negative T cells (n = 2 independent experiments and PBMC donors).

**D.** Upregulated ligand-receptor interactions identified among PBMC cell types after *P. falciparum*-stimulation using the *CellChat* package. Arrows point from ligands on sender cells to receptors on receiver cells and are colored by the sender cell. Weights of links are proportional to the interaction strength, and the strongest ligands and receptors are labeled.

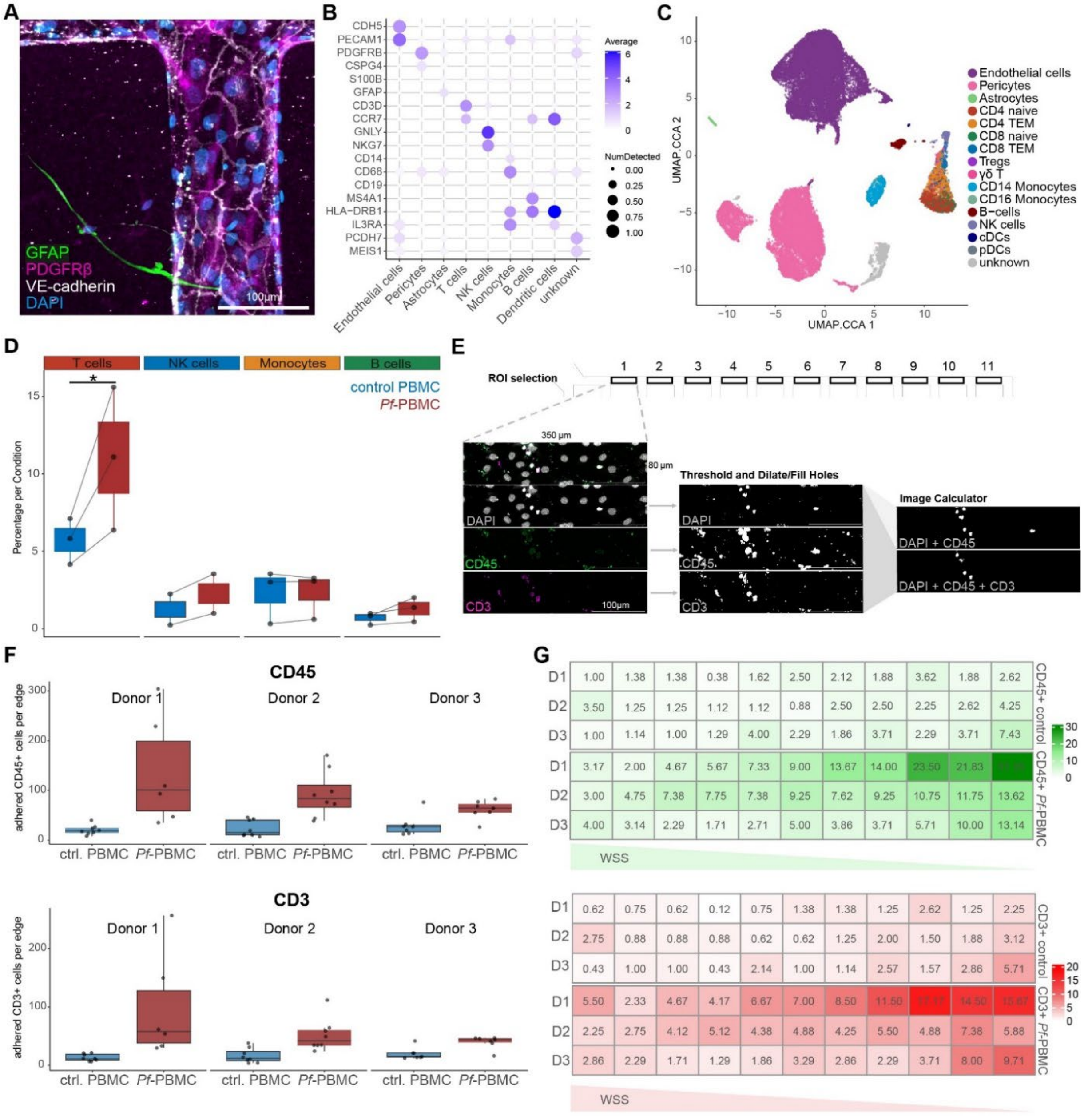


**Figure S2. Characterization of 3D-BBB model PBMC perfusion and immunofluorescence quantification of microvascular PBMC binding.**

**A.** Confocal microscopy image of the 3D-BBB model stained for main cell type markers glial fibrillary acidic protein (*GFAP*, astrocytes), platelet-derived growth factor receptor beta (*PDGFRβ*, pericytes), and VE-cadherin (endothelial cells).

**B.** Dot plot of canonical BBB and PBMC cell type markers in control and *Pf*-PBMC-perfused 3D-BBB devices.

**C.** UMAP visualization of control and *Pf*-PBMC-perfused 3D-BBB devices, colored by cell type annotation, including mapped subpopulations.

**D.** Box plot showing the proportion of each immune cell type adherent to the 3D-BBB model in control (blue) and *Pf*-PBMC-perfused (red) conditions of the scRNA-seq dataset. Each dot represents an scRNA-seq individual experiment. Asterisks labels statistically significant differences identified by *scCODA* compositional analysis (FDR < 0.05).
**E.** Image analysis pipeline for immunofluorescence quantification of microvascular PBMC binding, including maximum projection, thresholding, and overlay of binary masks (see Methods).
**F.** Quantification of adherent leukocytes stained for CD45 (top) and T cells stained for CD3 (bottom), split by PBMC donor and individual experiment. Each point represents one edge quantification coming from n=3 independent PBMC donors (two 3D-BBB devices per donor per condition).
**G.** Heatmap representation of quantification of adherent leukocytes stained for CD45 (top, green) and T cells stained for CD3 (bottom, red) along 3D-BBB device edges exhibiting a wall shear stress (WSS) gradient), split by PBMC donor and individual experiment. Numbers represent mean number of adherent cells per region of interest (ROI) (n = 6 3D-BBB devices and n = 3 PBMC donors per condition).

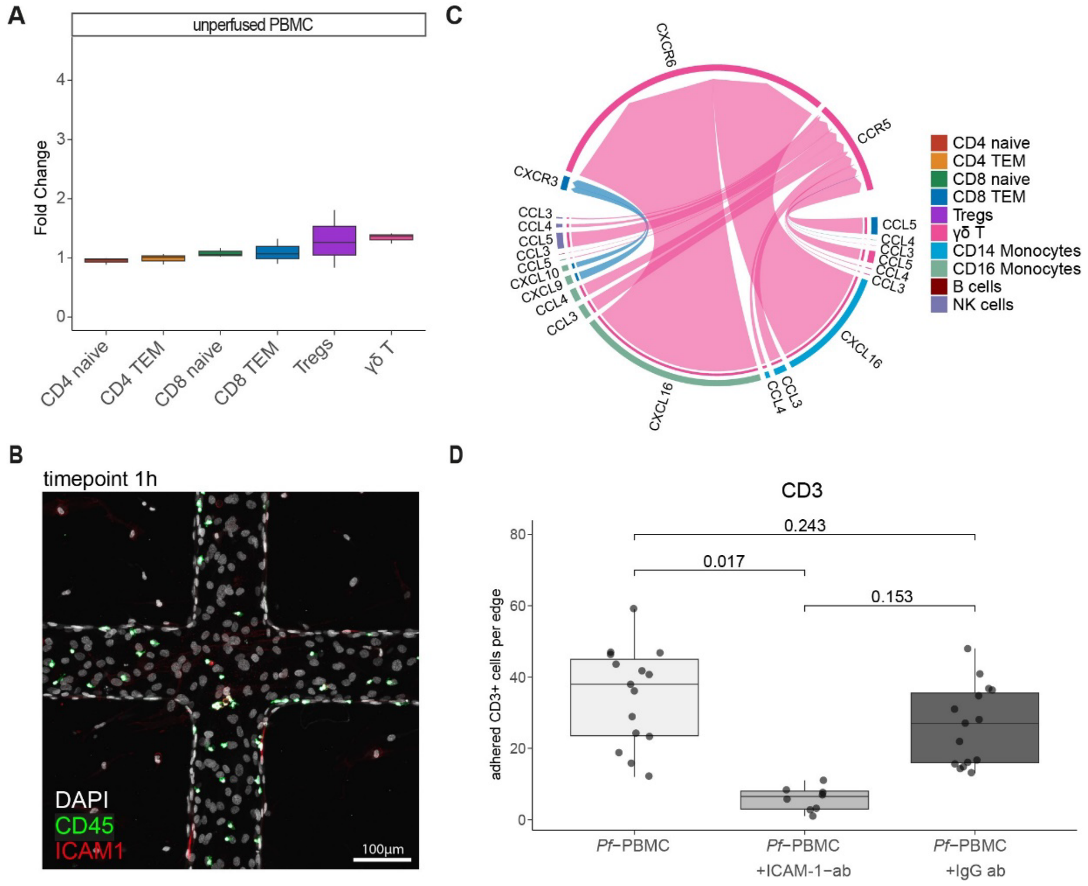


**Figure S3. *P. falciparum*-stimulated T cells show increased binding in 3D-BBB model.**
**A.** Fold change in the abundance of T cell subpopulations between control and *Pf*-PBMC (unperfused).
**B.** Confocal microscopy image of *Pf*-PBMC 3D-BBB model after 1-hour perfusion stained for ICAM-1 (red) and CD45 (green).
**C.** Chemokine receptor–ligand interactions (CXCL, CCL pathways) towards T cells in *Pf*-PBMC, identified using the *CellChat* package. Arrows point from sender cells to receiver cells and are colored by receiver cell. Arrow thickness is proportional to the interaction strength.
**D.** Number of adherent CD3-positive T cells per edge of the 3D-BBB model comparing control and *Pf*-PBMC perfused after 30-minute pre-treatment of the 3D-BBB model with a monoclonal ICAM-1-blocking antibody or an IgG isotype control antibody. Each point represents one edge quantification from 3D-BBB

devices perfused with *Pf*-PBMC (n=4 devices), ICAM-1 blocking antibody and *Pf*-PBMC (n=2 devices), or IgG control antibody and *Pf*-PBMC (n=4 devices), (two PBMC donors, two devices per donor per condition). Kruskal–Wallis test with Dunn's pairwise comparisons test and BH correction.

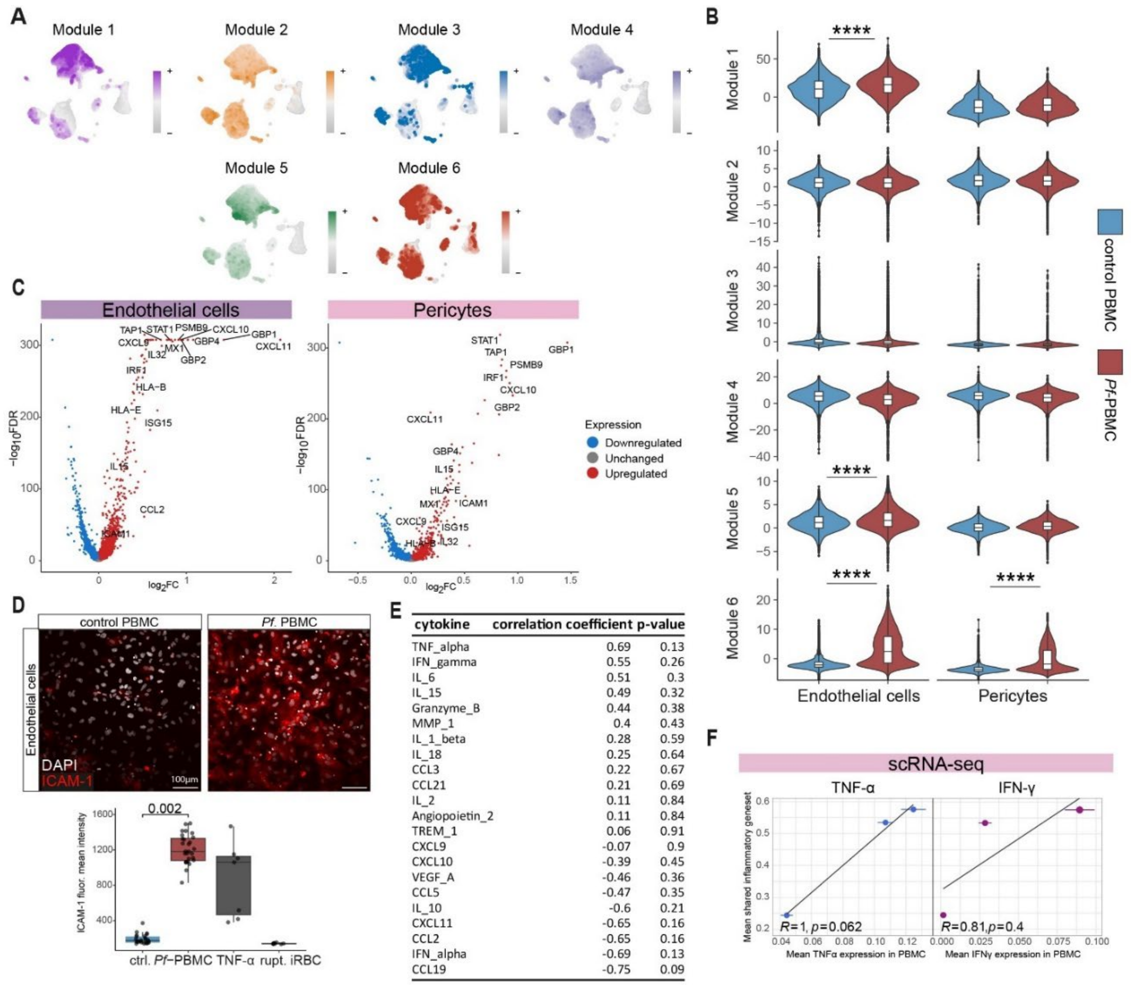


**Figure S4. Weighted gene network correlation analysis (WGCNA) and characterization of 3D-BBB inflammatory activation.**
**A.** UMAP plots showing eigengene expression of 6 transcriptional modules identified through WGCNA.
**B.** Violin plots showing the expression of 6 transcriptional modules in endothelial cells and pericytes of the 3D-BBB model exposed to control or *Pf*-PBMC. Asterisks mark significance with Mann Whitney U test and Bonferroni multiple testing correction (adjusted p-value < 0.05, expression in > 30% of cells, log2FC > 2).
**C.** Volcano plot of DE genes in endothelial cells and pericytes upon *Pf*-PBMC exposure, plotting the statistical significance (-log10 of the FDR) against the log2FC. Significantly up- or downregulated genes (FDR < 0.05) are marked in red or blue and selected upregulated genes are labeled.
**D.** Representative confocal maximum intensity z-projection showing ICAM-1 (red) labeling in endothelial cells after 10-hour incubation with control PBMC, *Pf*-PBMC, TNF-α, or ruptured iRBC in the same concentration as used for PBMC stimulation (0.18% parasitemia). Quantification of mean pixel fluorescence intensity is shown. Each dot represents on ROI from n=6 independent experiments and PBMC donors. Mann Whitney U test.

**E.** Correlation analysis between absolute cytokine levels (Luminex) in the supernatant of *Pf*-PBMC from n = 6 PBMC donors and induced ICAM-1 expression levels in endothelial cells. Pearson correlation coefficients and correlation p-values are listed.
**F.** Scatter plots showing the correlation between PBMC TNF-α and IFN-γ expression levels on the x-axis and induced inflammatory gene set expression (556 genes from Figure 4C) in endothelial cells on the y-axis. Each dot represents an individual scRNA-seq experiment. Pearson correlation coefficient (R) and p-values are indicated.

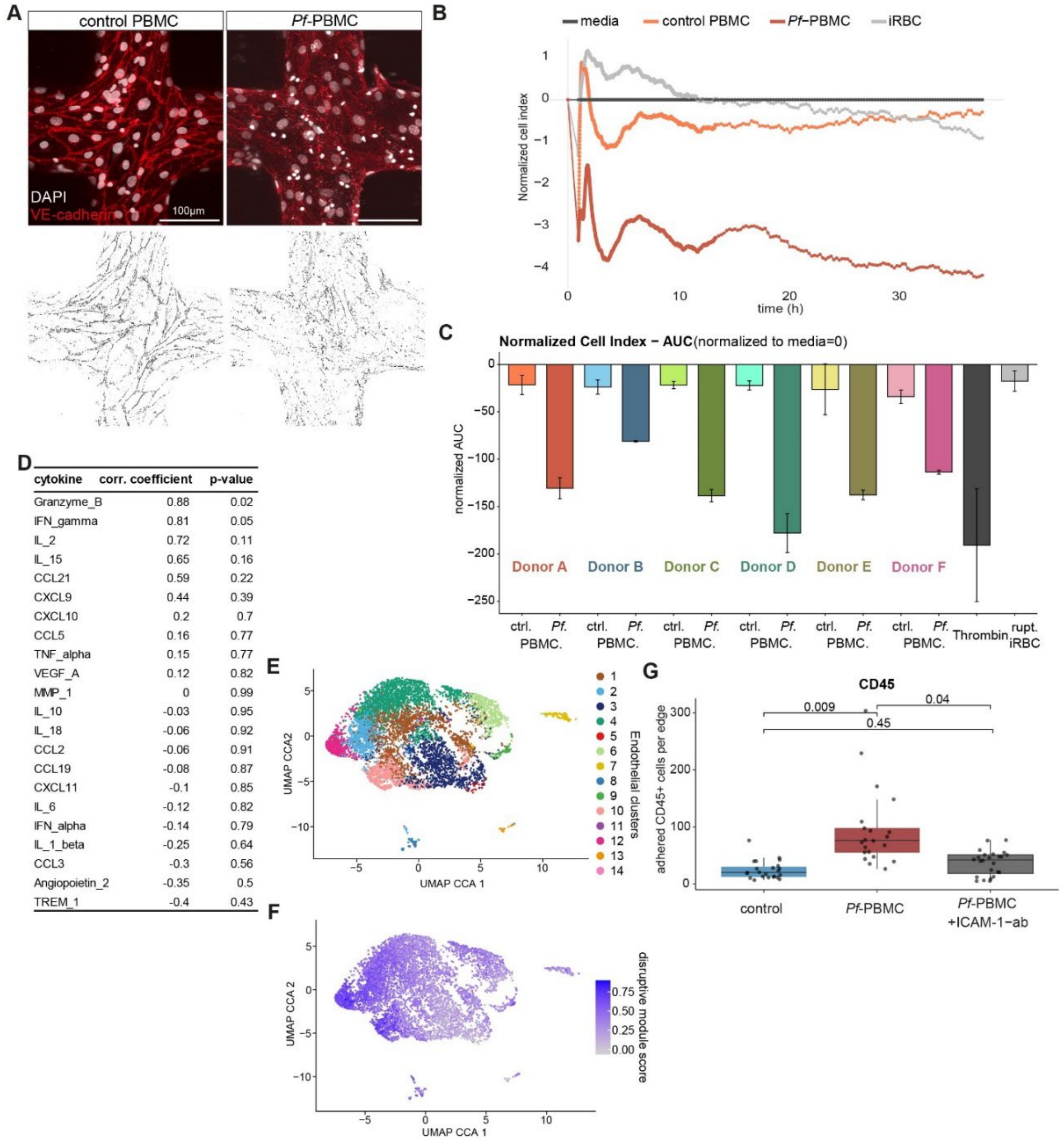


| cytokine | corr. coefficient | p-value |
|---|---|---|
| Granzyme_B | 0.88 | 0.02 |
| IFN_gamma | 0.81 | 0.05 |
| IL_2 | 0.72 | 0.11 |
| IL_15 | 0.65 | 0.16 |
| CCL21 | 0.59 | 0.22 |
| CXCL9 | 0.44 | 0.39 |
| CXCL10 | 0.2 | 0.7 |
| CCL5 | 0.16 | 0.77 |
| TNF_alpha | 0.15 | 0.77 |
| VEGF_A | 0.12 | 0.82 |
| MMP_1 | 0 | 0.99 |
| IL_10 | -0.03 | 0.95 |
| IL_18 | -0.06 | 0.92 |
| CCL2 | -0.06 | 0.91 |
| CCL19 | -0.08 | 0.87 |
| CXCL11 | -0.1 | 0.85 |
| IL_6 | -0.12 | 0.82 |
| IFN_alpha | -0.14 | 0.79 |
| IL_1_beta | -0.25 | 0.64 |
| CCL3 | -0.3 | 0.56 |
| Angiopoietin_2 | -0.35 | 0.5 |
| TREM_1 | -0.4 | 0.43 |

**Figure S5. Characterization of *Pf*-PBMC-induced BBB disruption.**
**A.** Example of VE-cadherin image analysis including Tophat filter, Otsu thresholding and measurement of VE-cadherin junctional area in the generated binary image.
**B.** Representative xCELLigence graph visualizing the cell index normalized to the media control on the y-axis over time (x-axis) for endothelial monolayers exposed to control or *Pf*-PBMC.

**C.** Area under the xCELLigence cell index curve (AUC) for endothelial monolayers exposed to control or *Pf*-PBMC from 6 different PBMC donors, normalized to media.
**D.** Correlation analysis between absolute cytokine levels (Luminex) in the supernatant of *Pf*-PBMC from n = 6 PBMC donors and induced ICAM-1 permeability increase in endothelial cells, expressed as absolute AUC. Pearson correlation coefficients and correlation p-values are listed.
**E.-F.** UMAP representation of *Pf.*-PBMC exposed endothelial cells, colored by unsupervised clustering (**E**) or *combined disruptive module score* (top hub genes from module 1, 5, 6) (**F**)
**G.** Number of adherent CD45-positive leukocytes per edge of the 3D-BBB model comparing control PBMC and *Pf*-PBMC (same data as Fig. 2F) with *Pf*-PBMC perfused after 30-minute pre-treatment of the 3D-BBB model with a monoclonal ICAM-1-blocking antibody. Each point represents one edge quantification coming from n=3 independent PBMC donors (two 3D-BBB devices per donor per condition), Kruskal–Wallis test with Dunn's pairwise comparisons test and BH correction

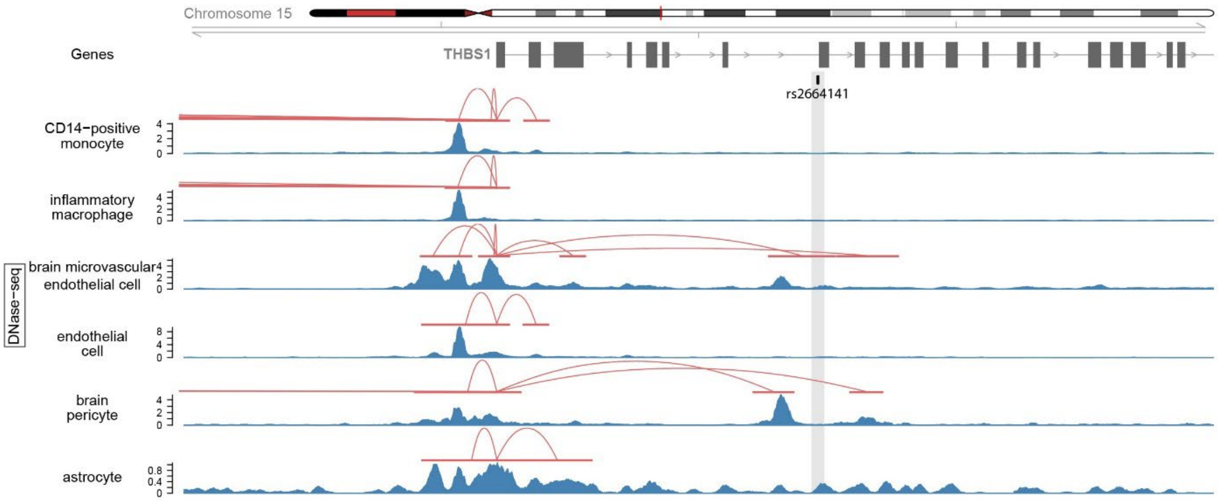


**Figure S6. Predicted cell type-specific enhancer-gene regulatory connections for *THBS1*.** Enhancer-gene regulatory connections of *THBS1* predicted by ENCODE-rE2G in cell types putatively involved in *THBS1* expression, visualized around an intronic SNP (rs2664141) found in a malaria-risk factor haplotype.
